# Evaluation of the safety, immunogenicity and efficacy of a new live-attenuated lumpy skin disease vaccine in India

**DOI:** 10.1101/2022.12.10.519851

**Authors:** Naveen Kumar, Sanjay Barua, Ram Kumar, Nitin Khandelwal, Amit Kumar, Assim Verma, Lokender Singh, Bhagraj Godara, Yogesh Chander, Thachamvally Riyesh, Deepak Kumar Sharma, Anubha Pathak, Sanjay Kumar, Ramesh Kumar Dedar, Vishal Mehta, Mitesh Gaur, Bhupendra Bhardwaj, Vithilesh Vyas, Sarjeet Chaudhary, Vijaypal Yadav, Adrish Bhati, Rakesh Kaul, Arif Bashir, Anjum Andrabi, Raja Wasim Yousuf, Abhimanyu Koul, Subhash Kachhawa, Amol Gurav, Siddharth Gautam, Hari Audh Tiwari, Madhurendu K. Gupta, Rajender Kumar, Jyoti Misri, Ashok Kumar, Ashok Kumar Mohanty, Sukdeb Nandi, Karam Pal Singh, Yash Pal, Triveni Dutt, Bhupendra N. Tripathi

**Affiliations:** National Centre for Veterinary Type Cultures, ICAR-National Research Centre on Equines, Hisar, India; Indian Veterinary Research Institute, Mukteswar, India; Department of Veterinary Microbiology, College of Veterinary and Animal Science, Navania, Vallabhnagar, Udaipur, India; Department of Animal Husbandry, Banswara, Rajasthan, India; Department of Veterinary Gynaecology and Obstetrics, College of Veterinary and Animal Science, Navania, Vallabhnagar, Udaipur, India; Department of Animal Husbandry, Udaipur, Rajasthan, India; Department of Animal Husbandry, Jodhpur, Rajasthan, India; Department of Animal Husbandry, Alwar, Rajasthan, India; Livestock Research station, Nohar, Rajasthan, India; Animal Husbandry Department, Jammu and Kashmir, India; Krishi Vigyan Kendra, ICAR-Central Arid Zone Research Institute, Jodhpur, India; Hasanand Gaushala, Vrindavan, Mathura, India; Department of Veterinary Pathology, Birsa Agricultural University, Ranchi, India; Animal Science Division, Indian Council of Agricultural Research, India; Centre for Animal Disease Research and Diagnosis, Indian Veterinary Research Institute, Izatnagar, India

**Keywords:** Lumpy skin disease, LSD/Ranchi, India, homologous live-attenuated LSD vaccine

## Abstract

Lumpy skin disease (LSD) was reported for the first time in India in 2019 and since then, it has become endemic. Since a homologous (LSD-virus based) vaccine was not available in the country, goatpox virus (GPV)-based heterologous vaccine was authorized for mass immunization against LSD in cattle. This study describes the evaluation of safety, immunogenicity and efficacy of a new live-attenuated LSD vaccine developed using an Indian field strain (LSDV/India/2019/Ranchi). The virus was attenuated by continuous passage (P=50) in Vero cells. The vaccine (50^th^ LSDV passage in Vero cells, named as Lumpi-ProVac^*Ind*^) did not induce any local or systemic reaction upon its experimental inoculation in calves (n=10). At day 30 post-vaccination (pv), the vaccinated animals were shown to develop antibody- and cell-mediated immune response and exhibited complete protection upon virulent LSDV challenge. We observed a minimum Neethling response (0.018% animals; 5 out of 26940 animals) of the vaccine in field trials among 26940 animals. There was no significant reduction in the milk yield in lactating animals (n=10108), besides there was no abortion or any other reproductive disorder in the pregnant animals (n=2889). Sero-conversion was observed in 85.18% animals in the field by day 30 pv.

## Introduction

Lumpy skin disease (LSD) is a transboundary animal viral disease which leads to considerable financial losses to the livestock industry. The disease is characterized by the development of skin nodules, fever, enlargement of lymph nodes, anorexia, depression, dysgalactia and emaciation which may eventually result in a sharp decline in the milk production, abortion in pregnant cows and infertility in bulls ^1^. LSD is a serious hazard to the food security of the people in the affected areas ^1, 2^. The World Organization for Animal Health (WOAH) categorizes LSD as a notifiable disease. The disease has remained restricted to Africa ever since its first occurrence^3 4^. Its first intercontinental spread was confirmed in Israel in 1989^5^. Since 2012, LSD has spread from Africa into several countries of Europe. It was first reported in the Asia and the Pacific region in 2019 from North West China, Bangladesh and India^6–8^. During 2020, LSD outbreaks continued to spread across many countries including Bhutan, Hong Kong, Myanmar, Nepal, Taiwan, Vietnam and Sri Lanka. In 2021, the disease was confirmed in Thailand, Vietnam and Malaysia, while Indonesia reported its first confirmed outbreak in March 2022.

LSD was reported for the first time in India in 2019^9^. Currently the country is facing the wrath of this deadly viral epidemic. Although, the mortality rate in LSD is usually considered low (<1%)^10^, however, it has been much higher during the current wave of LSD in India^1^. The loss of milk production in the affected cows has been reported to be between 26-50%^11^.

LSDV belongs to the genus *Capripoxvirus* within the family *Poxviridae*. LSDV genome is ~151 Kbp in length ^12^. Two other capripoxviruses, sheeppox virus (SPV) and goatpox virus (GPV) which causes severe disease in sheep and goats respectively, are also antigenically similar to LSDV^13^. Capripoxviruses are believed to provide cross protection within the genus; therefore SPV- or GPV-based vaccines have been used to provide cross protection against LSDV^13–17^. As an emergency measure, the policy makers in India authorized the use of goatpox vaccine (heterologous vaccine) against LSD in cattle in 2021 ^1, 18^. However, the cross protection issue has been controversial^14, 19, 20^. There are numerous examples wherein GPV/SPPV-based vaccines have been shown to induce partial protection against LSDV in cattle^13, 21–25^. These discrepancies in the use of heterologous vaccine prompted us to develop an indigenous homologous vaccine which confers solid immunity against LSD in cattle^13, 26^.

## Materials and Methods

### Ethics Statement

Vaccine efficacy experiments were conducted in calves at Indian Veterinary Research Institute (IVRI), Mukteswar, after obtaining due approval from the Committee for Purpose of Control and Supervision of Experiments on Animals (CPCSEA), Department of Animal Husbandry and Dairying, Ministry of Fisheries, Animal Husbandry and Dairying, Government of India (V-11011(13)/3/2022/CPCSEA-DADF, dated 10.03.2022).

The field trials were approved by National Research Centre on Equines, Indian Council of Agricultural Research, Hisar, India. A due consent was taken from the concerned farmer for inoculation of the experimental vaccine in cattle/buffaloes.

### Cells

Primary lamb testicle cells^27^ and African green monkey kidney (Vero) cells^28^ were available at National Centre for Veterinary Type Cultures (NCVTC), Hisar and grown in Dulbecco’s Modified Eagle’s Medium (DMEM) supplemented with antibiotics and 10-15% foetal calf serum.

### Viruses

LSDV/Cattle/India/2019/Ranchi-the field strain which was used to develop vaccine was isolated previously by our group in primary goat kidney cells from the skin nodules collected from cattle naturally infected with LSDV^9^. LSDV/Cattle/India/2021/Banswara, LSDV/Cattle/India/2022/Jalore and LSDV/Camel/India/2022/Bikaner were isolated by our group and are available at the National repository (NCVTC, Hisar India) with Accession Numbers of VTCCAVA 321, VTCCAVA 370 and VTCCAVA 371 respectively.

### Attenuation of LSDV in Vero cells

In order to attenuate, virus was sequentially passaged in Vero cells for up to 50 passages (P). For each passage, 500 μl inoculum of the virus having a titre of 10^6^ TCID_50_/ml from the previous passage was used to infect fresh Vero cells for 2 h, followed by washing with phosphate buffered saline (PBS) and addition of fresh growth medium. The virus was harvested when the cells exhibited ~75 % cytopathic effect (CPE).

### Whole genome sequencing

The P50 virus was completely sequenced at Next Generation Sequencing Platform (Clevergene Biocorp Pvt Ltd, Bangaluru, India). Viral DNA was extracted by DNeasy^®^ Blood & Tissue Kit (Qiagen, Hilden, Germany) as per the instructions of the manufacturer and sent to Clevergene Biocorp Pvt Ltd (Bengaluru, India) for whole genome sequencing. The complete nucleotide sequences of LSDV/India/2019/Ranchi/P0 and LSDV/India/2019/Ranchi/P50 were deposited to GenBank Accession Number of MW883897.1 and OK422494.1 respectively and the mutations were identified by using an online tool (https://www.ebi.ac.uk/Tools/msa/clustalo/).

### Virus neutralization assay

Serum samples were initially heated at 56°C for 30 min to inactivate the complements. Vero cells were grown in 96 well tissue culture plates till ~90 confluency. Two-fold serum dilutions (in 50 μl volume) were made in PBS and incubated with an equal volume of ~10^3^ TCID_50_ of LSDV for 1 h at 37 °C. Thereafter, virus-antibody mixture was used to infect Vero cells. The cells were observed daily for the appearance of CPE. Final reading was taken at 72 hours post-infection (hpi) for the determination of antibody titres.

### Vaccine preparation

The final vaccine preparation was carried out by diluting the original virus stock of LSDV/Ranchi/P50 (~10^7^ TCID_50_/ml) in sterile PBS to prepare aliquots of 10^4.5^ TCID_50_/ml (10X field dose) and 10^3.5^ TCID_50_/ml (1X field dose). The virus (vaccine) was tested for its sterility, wherein two ml of the final vaccine preparation was inoculated in 10 ml of Fluid thioglycollate medium (FTM) at 30°-35°C (for anaerobes) and Tryptic soy broth (TSB) at 20°-25°C (for aerobe /fungi) for 10 days. The sterility was ascertained by the absence of microbial contamination for up to 14 days. Besides, the final vaccine preparation was also tested for *Mycoplasma sps* and bovine viral diarrhoea virus (BVDV) by amplification of their respective gene segments in PCR.

### qRT-PCR

DNA was isolated from blood and nasal/fecal/ocular swab by DNeasy Blood & Tissue Kit (Qiagen, Hilden, Germany). The viral DNA was detected by TaqMan-probe-based real-time quantitative PCR as per the previously described method^29^. The primers flanked a conserved 151-bp region of LSD044 target region (forward primer-5’-CAA AAACAATCGTAACTAATCCA −3’; reverse primer-5’-TGGAGTTTTTATGTCATCGTC-3’). The probe (5’-6-FAM-TCGTCGTCGTTTAAAACTGA-QSY-3’) was labelled with 6-carboxy fluorescein (FAM), the reporter dye at the 5′-end and the QSY quencher at the 3′-end. Each 20-μL PCR reaction comprised 5 μL of DNA, 10 μL of 2 × Taq Man Universal PCR Master mix (Thermo Fisher Scientific, USA), 10 μM of each primer and 2.5 μM of the probe. PCR was conducted on a quantStudio 3 thermal cycler (Thermo Fisher Scientific, USA) with following Thermalcycler conditions-95°C for 5 min, followed by 40 cycles of 95°C for 15 s, and 60 °C for 60 s. Threshold cycle (Ct) values of ≤35 were considered as positive.

### Differentiation of LSDV vaccine and field (virulent) strains

LSDV ORF44 (*zdf4ln*) contains a conserved region which is present in all the LSDV strains but not in other capripoxviruses. This was exploited to specifically amplify LSDV genome (not SPV and GPV) by TaqMan real-time PCR as per the previously described method ^29^.

As compared to the field/virulent LSDV strains, most LSDV vaccine strains including Indian vaccine strain (LSDV/Ranchi/P50) contains a 12 bp insertion in its GPCR gene. This was exploited to specifically amplify the vaccine strain (but not field strains) by TaqMan real-time PCR as per the previously described method^30^.

### Preparation of challenge virus

Skin nodules were collected from a naturally LSDV infected cattle. A 10% suspension of skin nodules was used to infect the primary goat kidney cells. At 4 days following infection, the cells were freeze-thawed twice to harvest the virus (P1). The P1 virus was bulk cultured in primary lamb testicle cells and concentrated 50-times by ultracentrifugation. The final virus preparation (P2) had a virus titer of 10^5.5^ TCID_50_ (Ct value of 19.6 in qRT-PCR).

### Safety and efficacy

The safety and efficacy was conducted as per WOAH Terrestrial Manual 2021^31^. LSDV seronegative male calves, aged 6-9 months, were included in the study. Out of the 15 calves used to evaluate the safety and efficacy of the vaccine, two animals were inoculated with 10^4.5^ TCID_50_, while eight were inoculated with 10^3.5^ TCID_50_ and 5 were kept as unvaccinated control.

Animals were monitored for any clinical signs and rectal temperatures were recorded. On day 30 after vaccination, Thereafter, the animals were challenged with virulent virus by intraveneous (2 ml) and intradermal (0.25 ml at four sites on the flank region) routes on day 30 pv. The clinical response was recorded for 17 days.

### Safety in the field animals

A total of 26940 animals (cattle and buffaloes) of all age groups including lactating (n=10108) and pregnant (2889) animals were vaccinated with a recommended field dose (10^3.5^) of the vaccine by subcutaneous route. All the animals were observed for development of any local or systemic reactions. Body temperature was recorded in selected farms. Total daily milk production in lactating animals and reproductive disorders (abortion) in pregnant animals were also recorded.

### Cell proliferation assay

Peripheral blood mononuclear cells (PBMCs) were isolated from blood by HISTOPAQUE^®^-1077 (Sigma, Steinheim, Germany) as per the instructions of the manufacturer and resuspended in RPMI (Sigma, St Luis, USA) supplemented with 15% FBS. The cells were cultured in 96-well tissue culture plates at concentrations of 2×10^6^ cells/ml in 100 μl volume and stimulated with either 10 μg of UV-inactivated LSDV antigen- or Concavalin A (positive control). A negative control was made up of unstimulated PBMCs in cell culture medium only. Cultures were incubated at 37°C in 5% CO_2_ for 72 h. Five milligrams of MTT was dissolved into 1 ml of PBS, with 50 μl added onto each well containing cells and incubated at 37°C for 4 h. Finally 100 μl of DMSO was added to dissolve the formazan crystals. The absorbance (OD) was measured at 570 nm and relative increase in cell numbers in LSDV-Antigen stimulated over un-stimulated wells was determined.

### Measurement of IFN-ϒ

The levels of IFN-ϒ in naïve, vaccinated, vaccinated-challenged and unvaccinated-challenged animal were measured by Bovine IFN-ϒ-ELISA kit (Invitrogen, Frederick, USA) as per the instructions of the manufacturer.

### Statistical analysis

Effect of vaccination on different biological parameters in vaccinated and control animals was compared using Student’s t-test

## Results

### Genetic signatures of vaccine virus (LSDV/India/2019/Ranchi/P50)

Like other poxviruses, *Capripoxvirus* genome is quite large (151 Kbp) and undergoes extensive mutations during cell culture passage. Besides several point- and frameshift mutations, insertions and deletions of DNA segments can also be observed during the evolution of *Capripoxviruses*^32, 33^. We compared the whole genome sequences of LSDV/Ranchi/2019 at P0 (field strain, GenBank Accession Number MW883897) and at P50 (50^th^ Vero passage, GenBank Accession Number OK422494). As compared to the LSDV/P0, the major mutations in LSDV/P50 were observed in genes encoding Ankyrin repeat proteins, Kelch-like proteins, EEV membrane phosphoglycoprotein and DNA-dependent RNA polymerase. Nevertheless, these viral genes have also been shown to be disrupted in other live-attenuated vaccine candidates of capripoxviruses (LSDV, SPV and GPV)^33^. A twelve bps insertion in G-protein coupled chemokine receptor (GPCR)-the signature mutation of most of the *Capripoxvirus* vaccine strains^30^ was also present in LSDV/Ranchi/P50. Interestingly, in contrast to other existing field and vaccine strains, a unique deletion of 801 bp in inverted terminal repeat region (ITR) was also observed in LSDV/Ranchi/P50.

### Preparation of vaccine

The LSDV/P50 produced CPE within 2-3 days (depending on MOI used to infect). The optimum yield (10^7^ TCID_50_/ml) of LSDV/P50 in Vero cells was obtained at MOI of 0.1 and at a virus harvest time of 72 h **(Fig. 1).** Besides sterility, the virus stock was also tested negative for extraneous agents viz; *Mycoplasma sps* and Bovine viral diarrhoea virus. The experimental trials were conducted with frozen stock of LSDV/Ranchi/P50. For the field trials, freeze-dried virus (vaccine) was used. To prepare freeze dried stock (50 doses pack size), the original stock of LSDV/Ranchi/P50 (10^7^ TCID_50_/ml) was diluted 100-times in PBS and 1 ml of it was dispensed into each vial (10^5.5^ TCID_50_/ml per 50 doses). Since freeze-drying resulted in a loss of ~5-fold virus titre, 5-times higher concentration (10^5.5^ TCID_50_/ml per 50 doses; 10^4^ TCID_50_/ml per dose) was considered for freeze-drying. The freeze-dried virus had a virus titer of 10^3.5^ TCID50. It was sterile and free from extraneous agents and named as Lumpi-ProVac^Ind^.

**Fig. 1:**
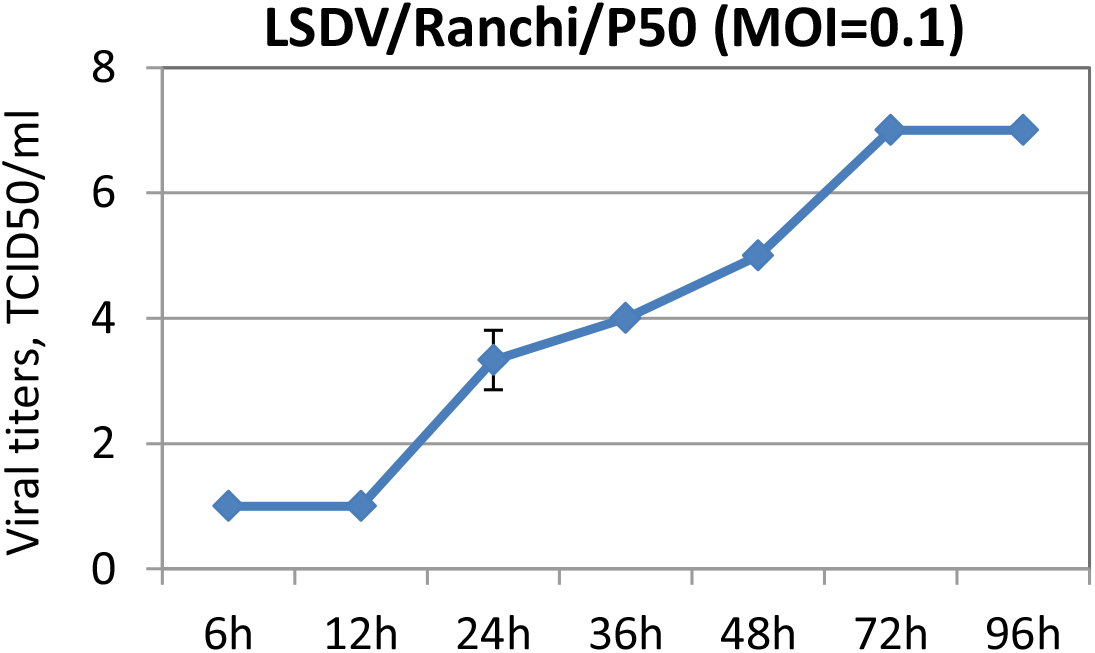
Growth characteristics of LSDV/Ranchi/P50. Vero cells, in triplicates, were infected with LSDV/Ranchi/P50 at MOI of 0.1 for 2 h, followed by washing with PBS and addition of fresh DMEM. Supernatant was collected from the infected cells at indicated time points and quantified by determination of TCID_50_ in Vero cells. Values are means ± SD and representative of the result of at least 3 independent experiments.

### Safety (Experimental trial)

Vaccinated calves did not develop any local or systemic reaction. Rise in temperature was recorded in a total of three animals viz; day 1 post-vaccination (pv) (IVRI 1529), day 3 pv (IVRI 1541 and IVRI 1529) day 12 pv (IVRI 1519) and day 15 pv (IVRI 1529) **(Supplementary Table 1).** Viral genome was detected in some of the vaccinated animals at day 3 pv **(Table 1),** however, the virus could not be isolated in any of the animals. Nasal, ocular and fecal shedding was not reported in any of the immunized animals up to day 30 pv **(Table 1).** All the immunized animals remained apparently healthy with a normal feed intake without exhibiting any untoward reaction. Various blood parameters of vaccinated (n=10) and unvaccinated (n=5) animals were also comparable **(Supplementary Fig S1).**

**Table 1:**
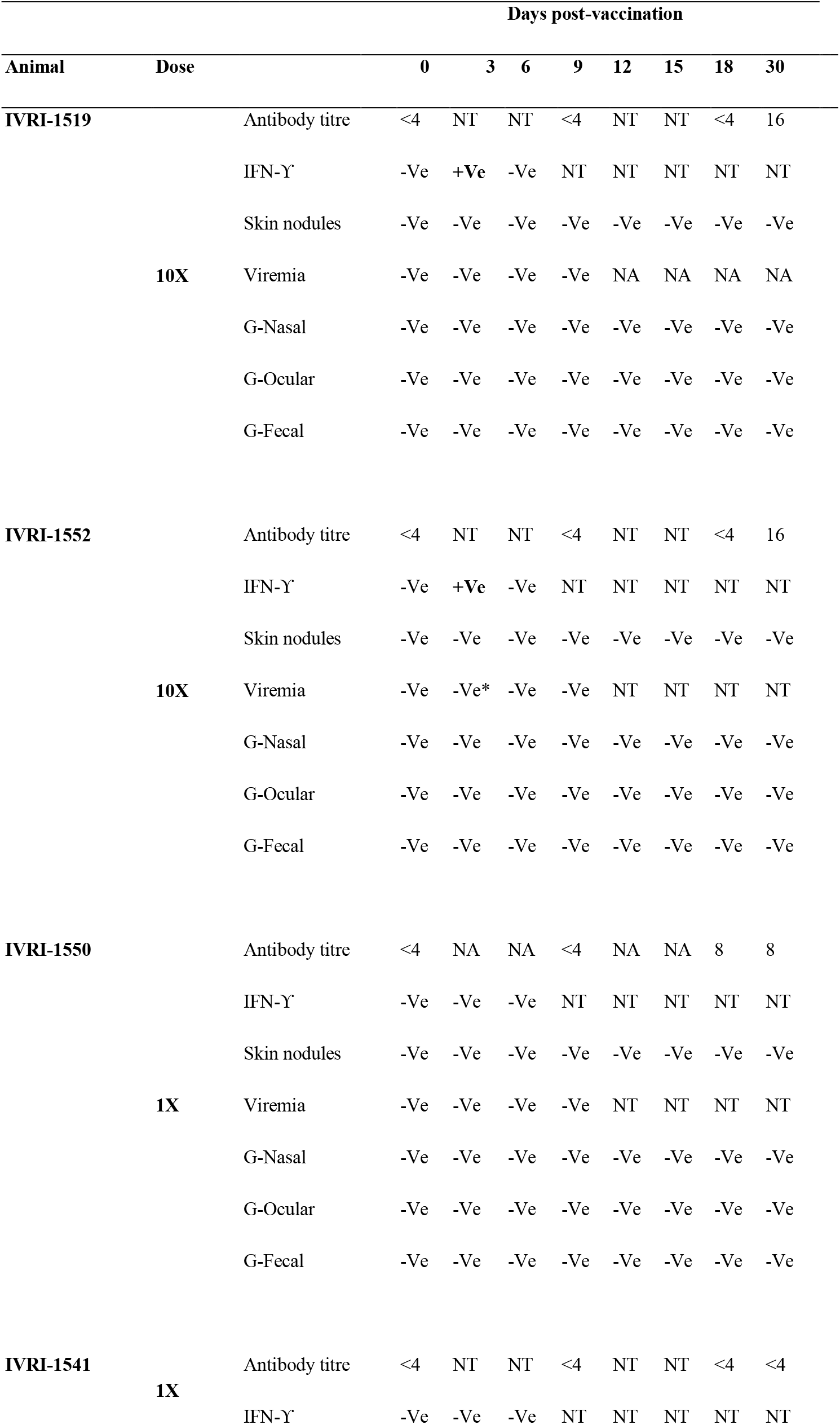
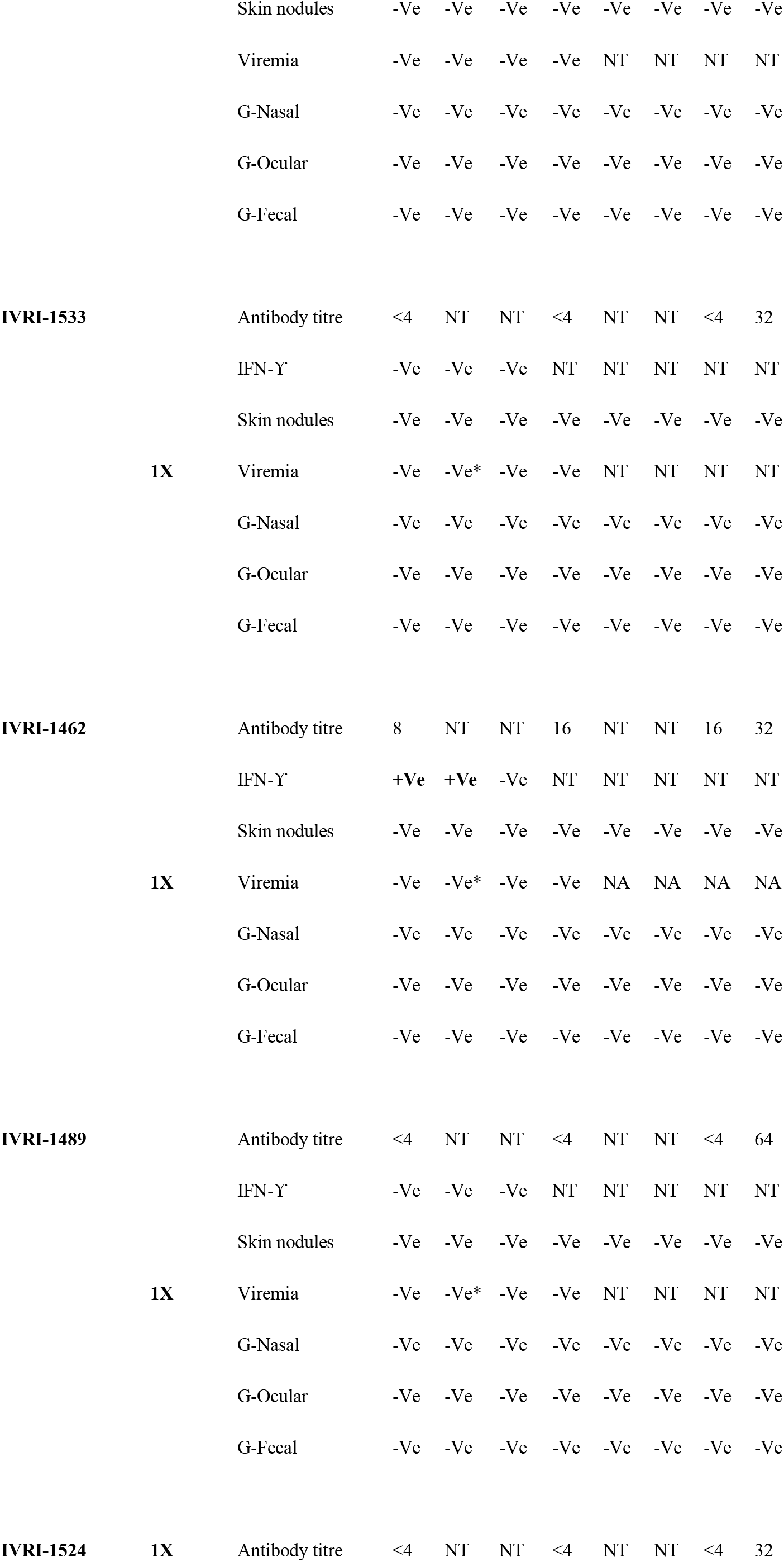
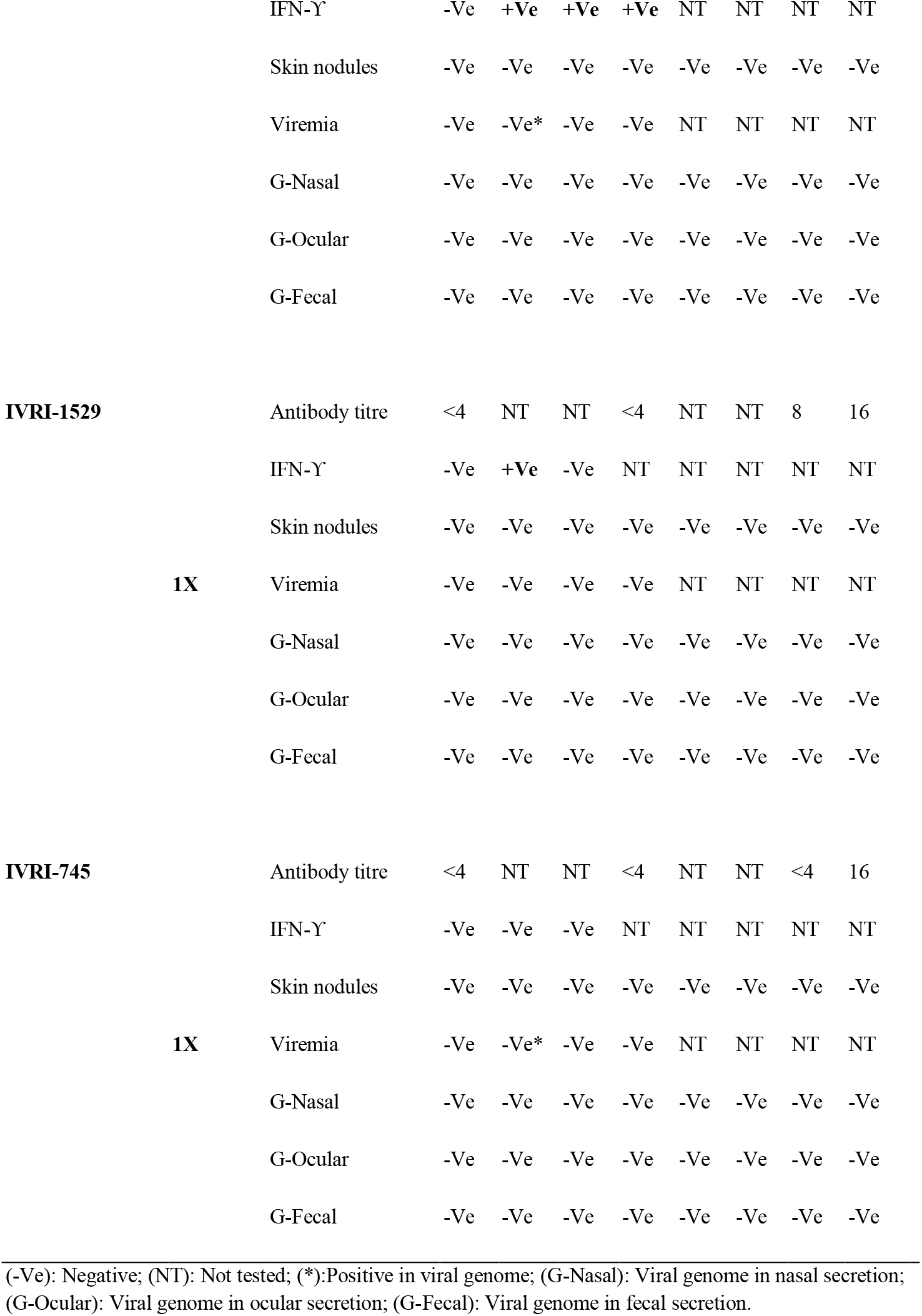
Observation of various safety parameters following vaccination with Lumpi-ProVacInd (Experimental trial)

### Safety in the field animals

A total of 26940 animals (26527 cattle and 413 buffaloes) across six different states in India **(Fig. 2a, 2b)** were included in the study. A single field dose of the vaccine (10^3.5^ TCID_50_) was found to be safe in cattle and buffaloes of all age groups including calves, pregnant/lactating animals and bulls. Fever was not observed in the field animals, although daily body temperature was noted in selected farms **(Supplementary Table 2 and 3).** Very mild swelling response (local site reaction which appeared at day 3 pv and subsided within 2 days without any specific intervention/treatment) was observed in 5 out of the 26940 vaccinated animals **(Table 2, Fig 2c).** Generalized skin nodules were not observed in any of the vaccinated animals. Likewise, abortion or any other reproductive disorders were also not observed in any of the 2889 pregnant (3 to 9 months of pregnancy) animals that received the vaccine **(Table 2).** Out of the 102 farms/units included in the study, a slight (non-significant) drop in milk production was recorded only in 4 farms/units (235 litres out of a total of 536024 litres in 8 days) **(Table 2, Fig. 2c)**. However this drop in milk production was temporary and was regained within 1-8 days pv **(Table 2).** All the vaccinated animals remained apparently healthy following vaccination without any significant alteration in feed/water intake.

**Fig. 2:**
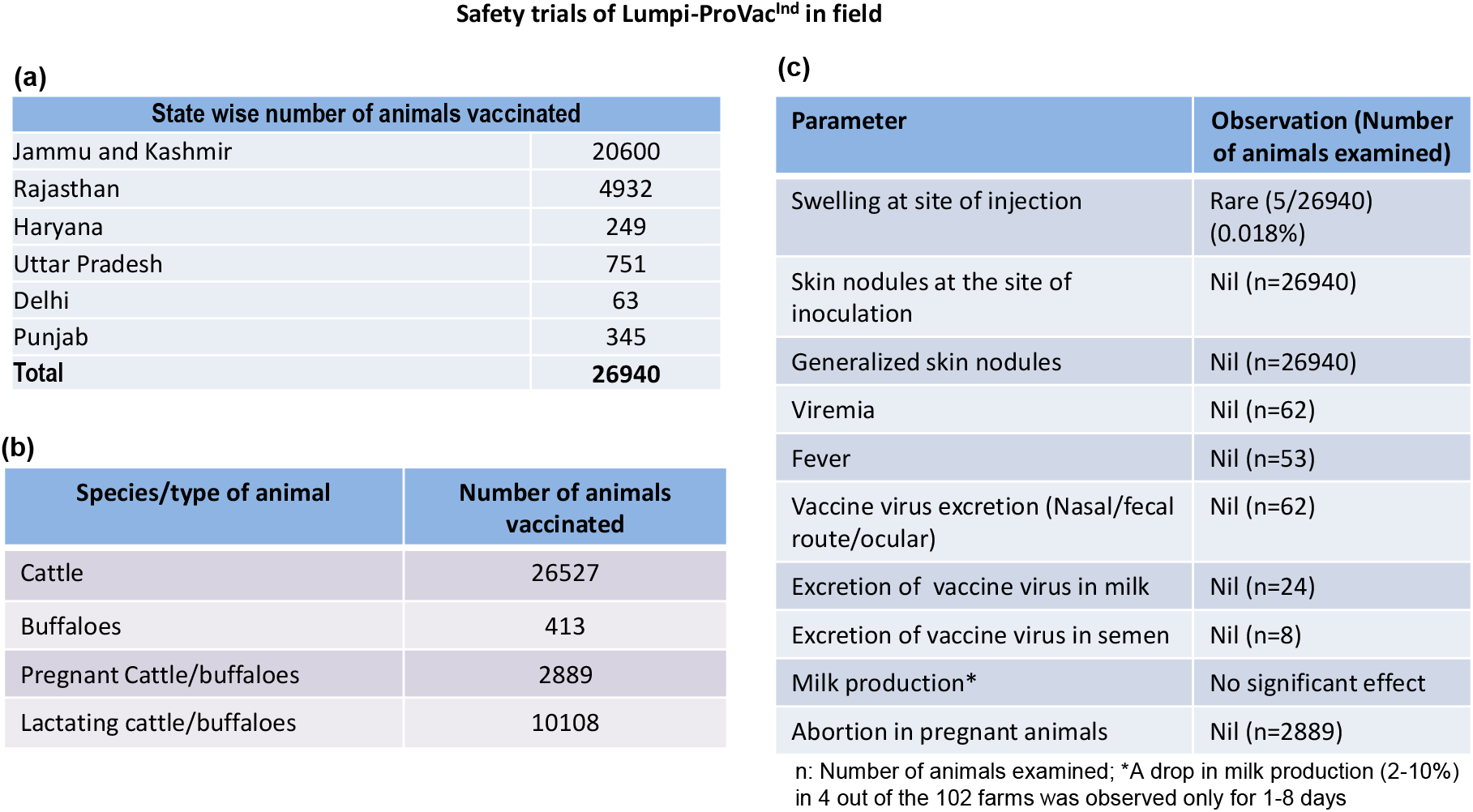
Safety of Lumpi-ProVac^Ind^ in field animals. A total of 26940 animals across six Indian states **(a)** comprising of 26527 cattle, 413 buffaloes (2889 pregnant cattle/buffaloes and 10108 lactating buffaloes) **(b)** were included in the study. All the animals were injected with 1 ml of Lumpi-ProVac^Ind^ (containing 10^3.5^ TCID_50_/dose) by subcutaneous route and monitored for swelling/skin nodules at the site of injection and generalized skin nodules **(c).** Besides, selected animals were also monitored for fever, viremia, virus excretion (nasal/fecal/ocular secretions and in milk/semen), milk production and reproductive disorders (abortion).

**Table 2:**
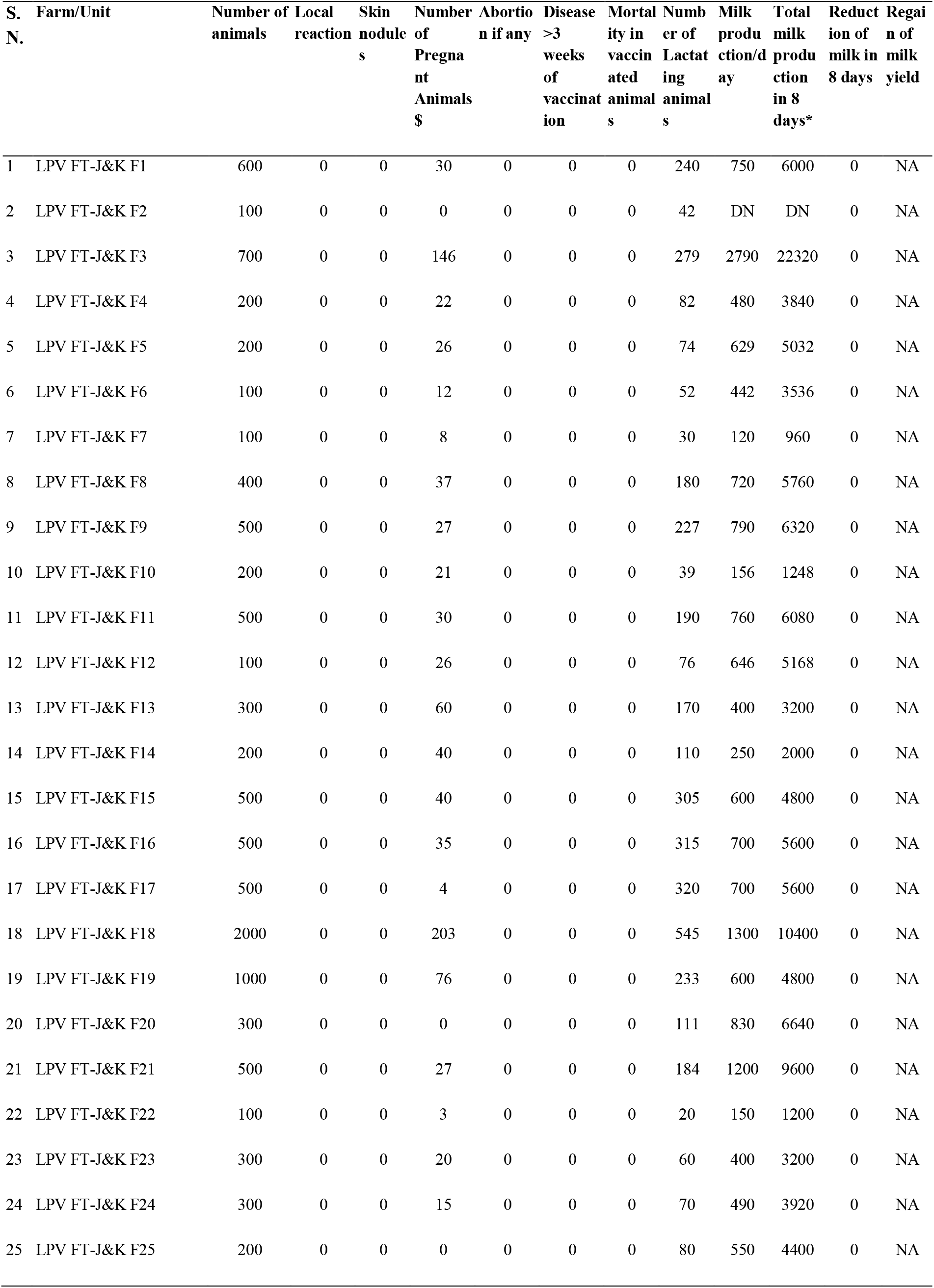
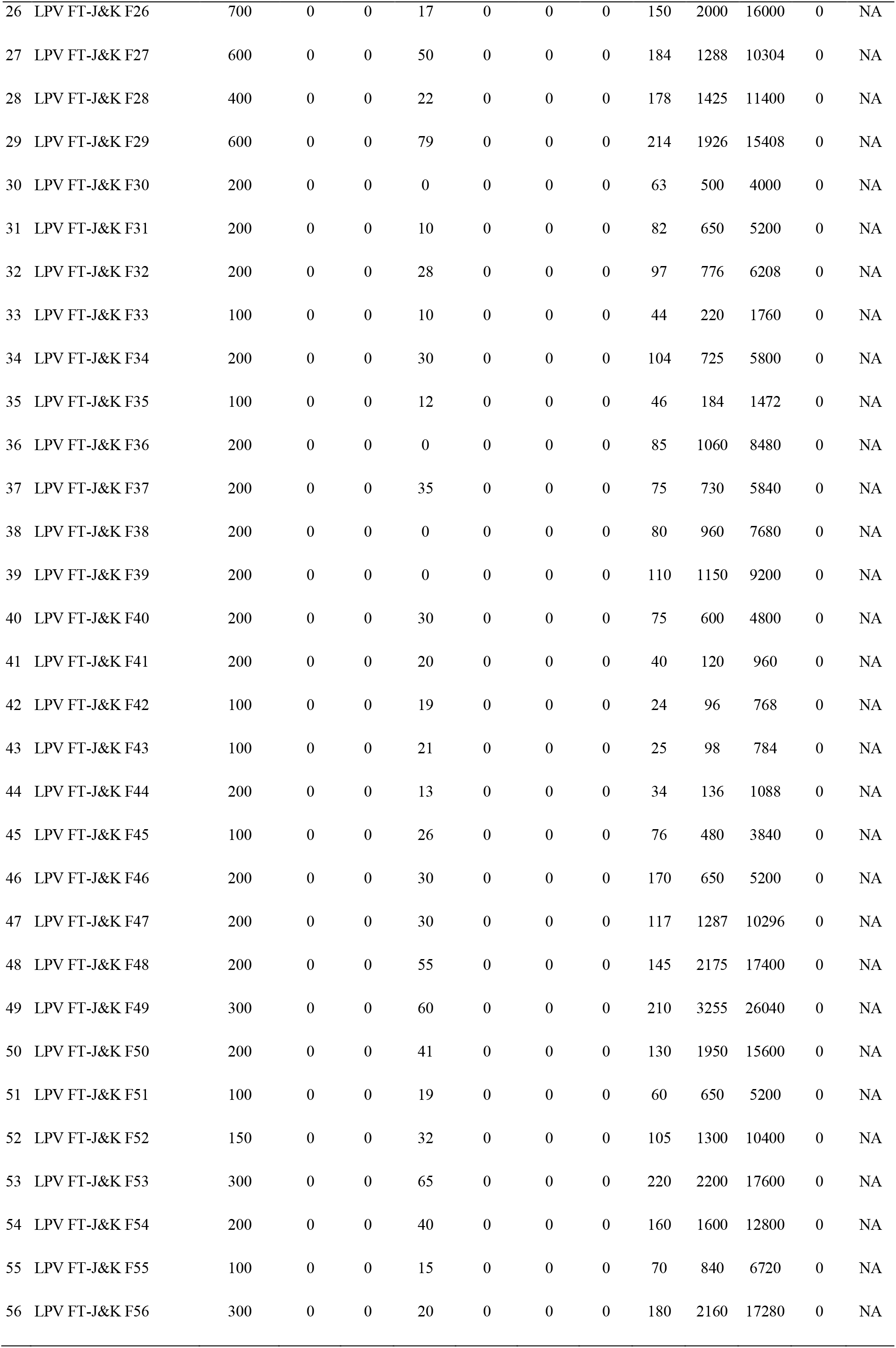
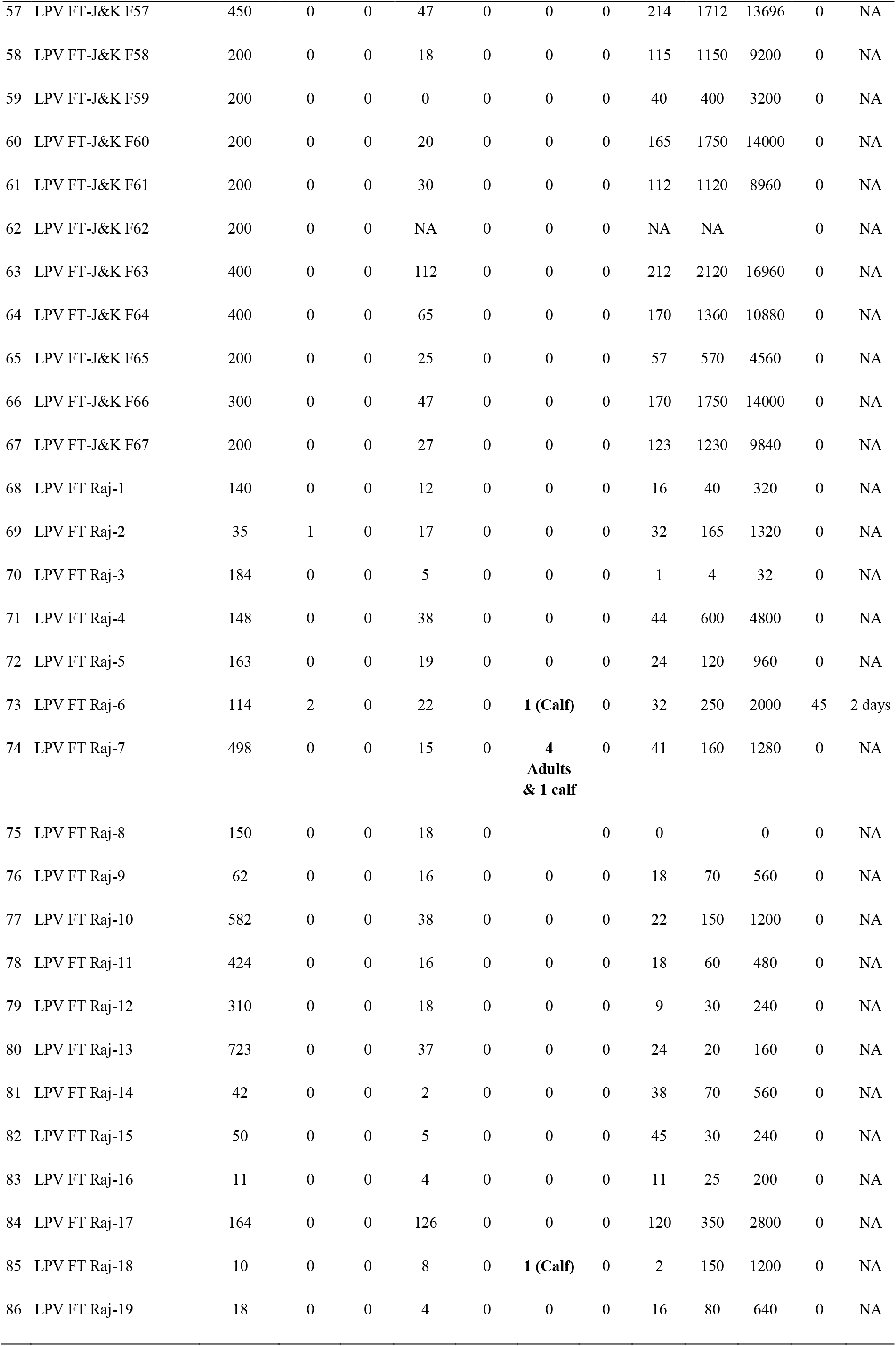
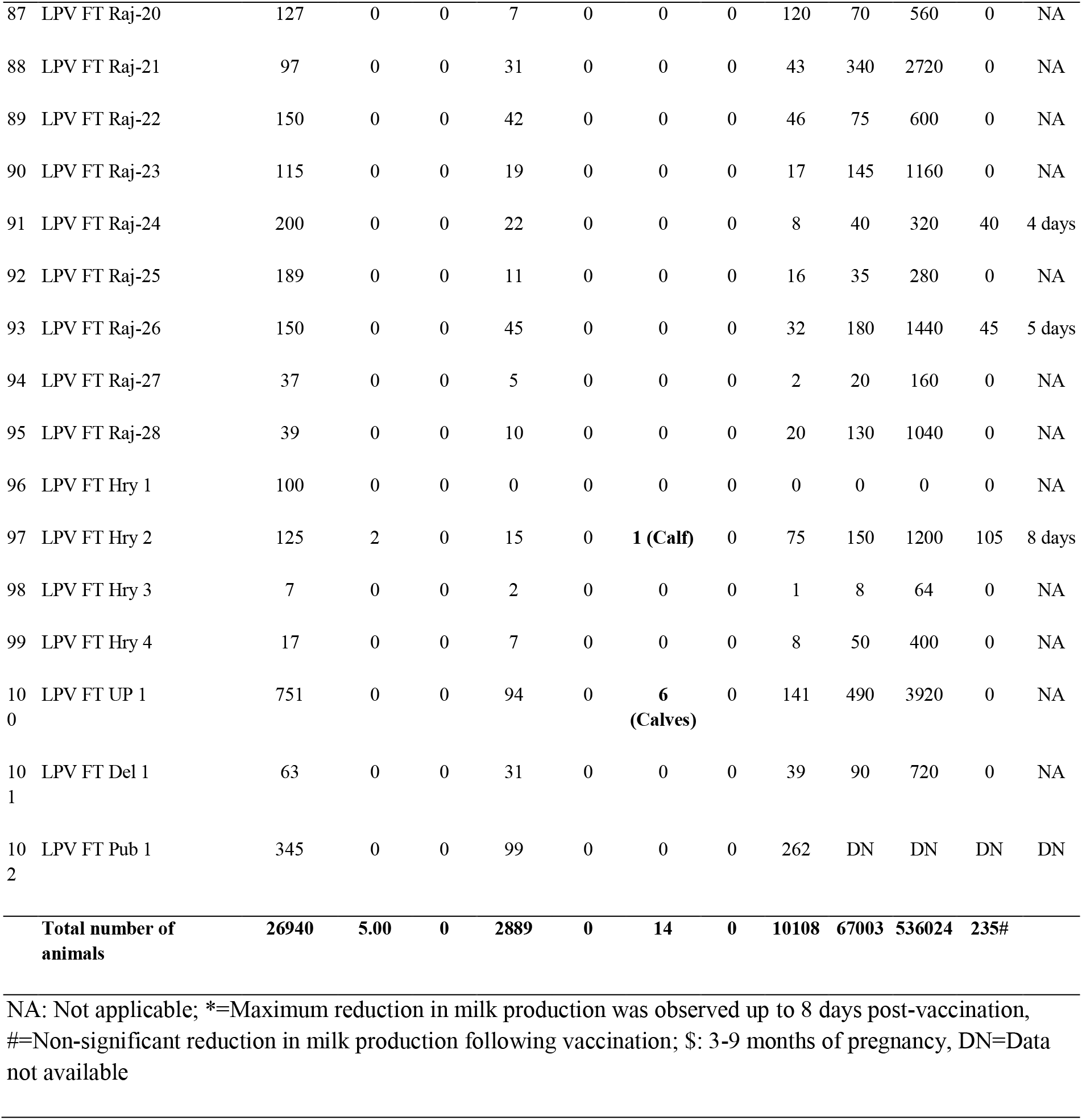
Safety and efficacy of Lumpi-ProVacInd in field animals.

### Immunogenicity

Serology of vaccinated animals, evaluated by virus neutralization test (VNT) revealed 90% positive animals at day 30 pv (experimental trial) **(Table 1, Fig. 3a)**. Antibodies were observed starting from day 18 pv. At 30 day pv, the antibody titter ranged between 8-64 **(Table 1)**. One of the cattle (IVRI 1541) did not reveal detectable amount of antibodies up to day 30 pv **(Table 1).** Although, prior to the start of experiment, animals were screened for seronegativity (free from antibodies against capripoxviruses) but one animal (IVRI 1462) had pre-existing antibodies at day 0 pv. In field trials, seroconversion was detected in 85.15% animals (n=648) (**Fig 3a).**

**Fig. 3:**
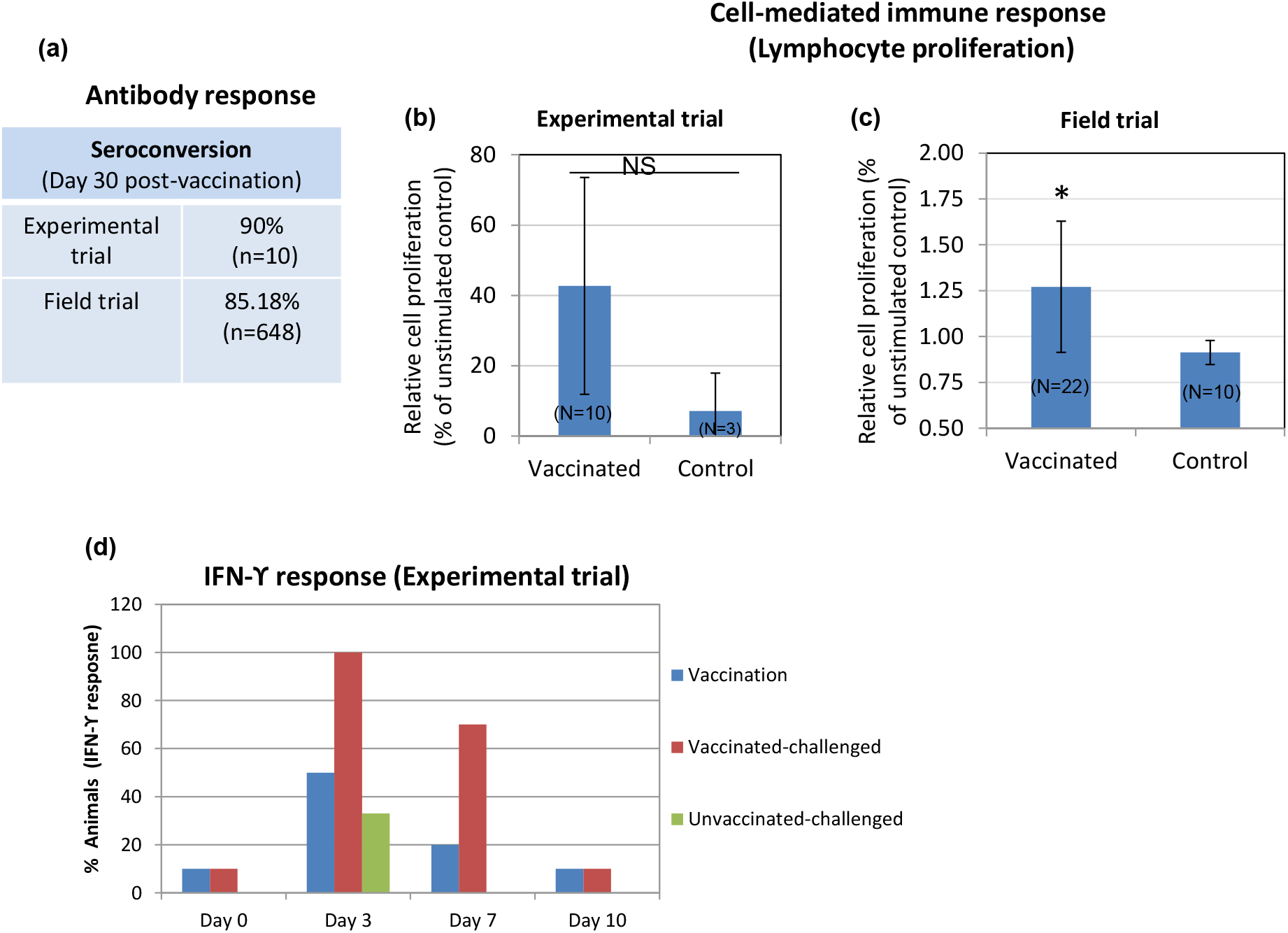
Immunogenicity of Lumpi-ProVac^Ind^. Immunogenicity was evaluated in experimental animals (n=10) and selected field animals (n=22). **(a) Antibody titers.** Percentage of animals (experimental and field trials) that revealed detectable anti-LSDV antibodies in serum by virus neutralization assay is shown. **(b) Lymphocyte proliferation assay (experimental trial).** PBMCs were separated from the blood collected from vaccinated (n=10) or unvaccinated (n=3) calves at day 30 pv. PBMCs were cultured in RPMI and stimulated with UV-inactivated LSDV. Relative proliferation of lymphocyte from vaccinated as compared to unvaccinated animals is shown. **(c) Lymphocyte proliferation assay (field trial).** PBMCs were separated from the blood collected from vaccinated (n=22) or unvaccinated (n=10) animals (all age groups) at day 30 pv and stimulated with UV-inactivated LSDV. Relative proliferation of lymphocyte from vaccinated as compared to the unvaccinated animals is shown. **(d) IFN-ϒ response.** Serum separated from the vaccinated, vaccinated-challenged and unvaccinated-challenged animals at indicated times post-LSDV exposure was examined for determination of IFN-ϒ by using Bovine IFN-ϒ-ELISA kit. Percentage of animals that exhibited IFN-ϒ response at indicated times points is shown.

We also evaluated the cell-mediated immune response by cell proliferation assay and detection of IFN-ϒ in serum. At day 30 pv, as compared to the unvaccinated controls, PBMCs from 60% of the immunized animals showed a significant cell proliferation response following stimulation with UV-inactivated LSDV antigen **(Table 3),** although, due to large animal-to-animal variation, the overall relative difference in cell proliferation was non-significant **(Fig. 3b).** However, in field trials, a significant difference in cell proliferative response was observed between vaccinated (n=22) and unvaccinated (n=10) animals **(Fig. 3c).**

**Table 3:**
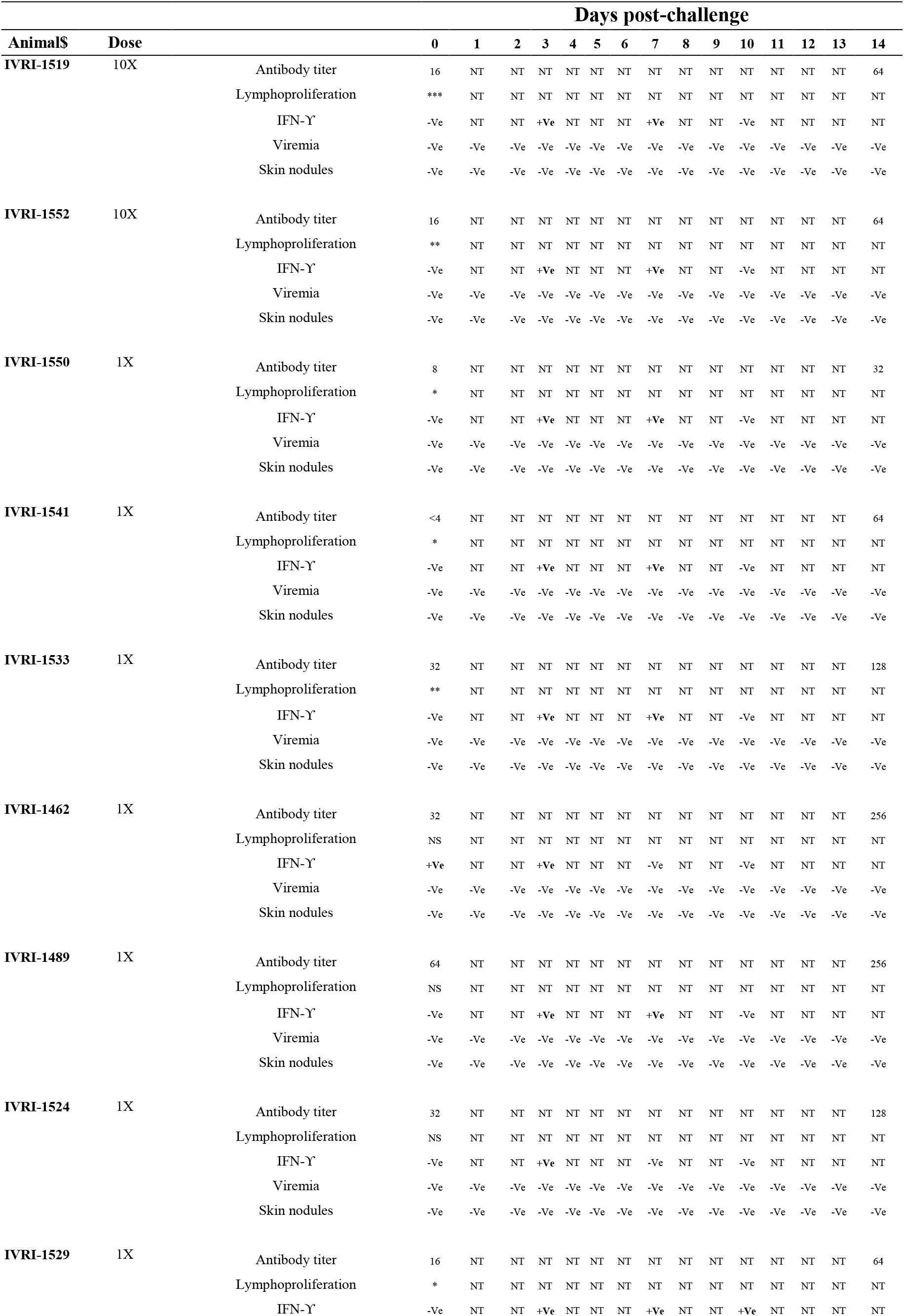
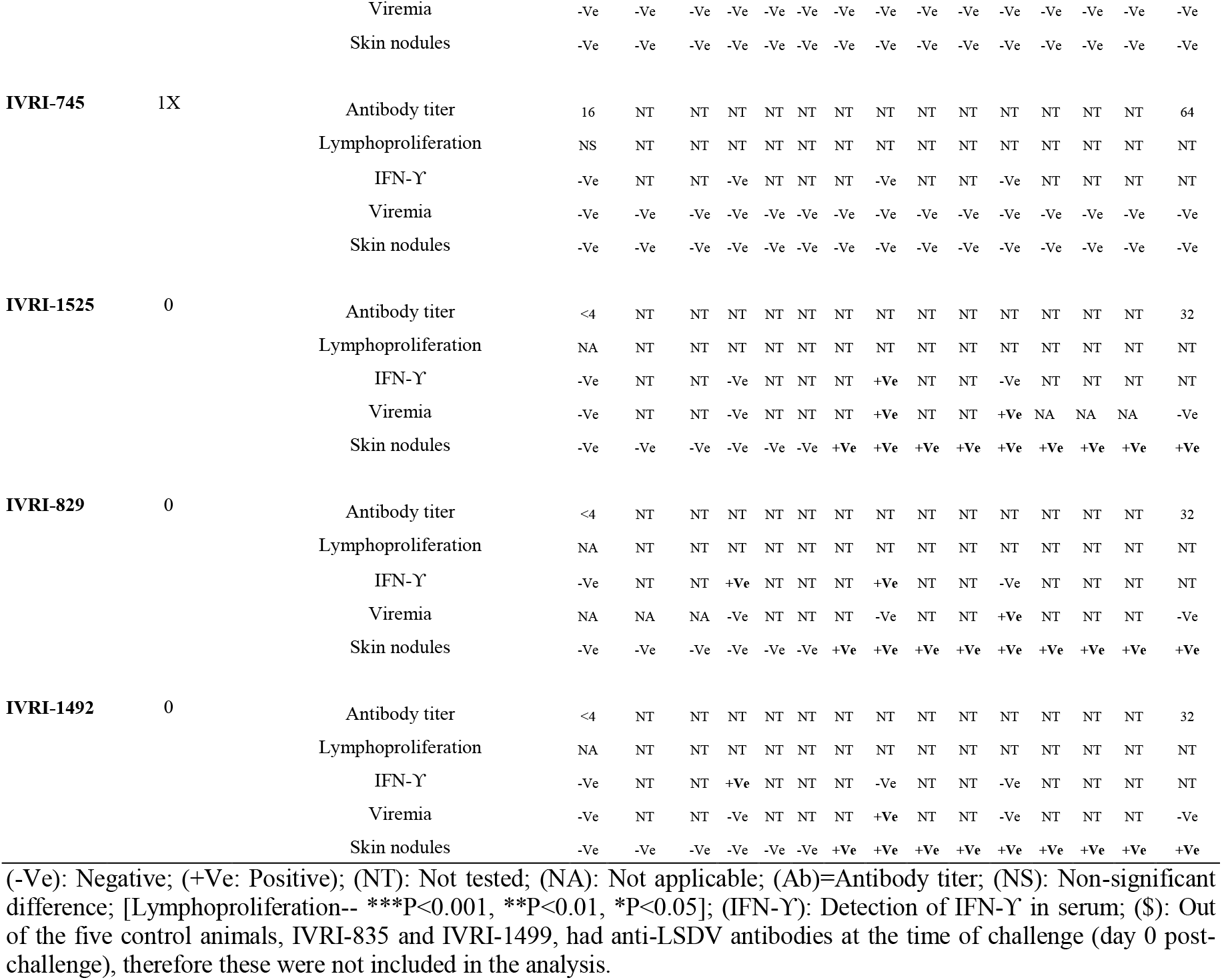
Efficacy of Lumpi-ProVacInd in cattle (Experimental challenge)

Following vaccination, the IFN-ϒ response was observed in 10%, 50%, 10% and 0% animals on day 0, day 3 day 7 and day 10 pv respectively **(Table 3, Fig. 3d)**. The challenge of vaccinated animals (n=10) showed IFN-ϒ response in 0%, 100%, 70% and 10% animals on day 0, day 3, day 7 and day 10 post-challenge (pc) respectively **(Table 3, Fig. 3d)**. Among the unvaccinated-challenge group (n=3), the IFN-ϒ response was observed in 0%, 66%, 33% and 0% animals on day 0, day 3, day 7 and day 10 pc respectively **(Table 3, Fig. 3d).** To summarise, the highest IFN-ϒ response was observed at day 3 following exposure with LSDV, irrespective of vaccination or challenge.

### Efficacy (Experimental trial)

All the immunized animals (n=10) along with the unvaccinated controls (n=5) were challenged with the virulent LSDV on day 30 pv. All the control animals (n=3 as 2 of the 5 control animals showed anti-LSDV antibodies in serum at day 0 pc and therefore were not included in the analysis) developed fever between 5-9 days pc which lasted for 1-4 days **(Supplementary Table 4)**. Localized skin nodules began appearing in all the unvaccinated control animals from day 5 to 6 pc **(Table 3).** The size of the skin nodules progressively increased from ~600 mm^2^ (day 6 pc) to 5000 mm^2^ (day 16 pc) before stabilizing at day 17 pc **(Fig. 4a** and **4b).** Skin nodules were observed at all four sites in all the inoculated animals. Few generalized skin nodules could also be observed in the unvaccinated control animals, although large numbers of skin nodules, as seen in natural infection were not apparent. In addition, viremia was also observed in all the unvaccinated-challenged animals between day 7-10 pc **(Table 3)**. Besides, the unvaccinated control animals were also found to be anorectic and depressed on day 6-8 pc.

**Fig. 4.**
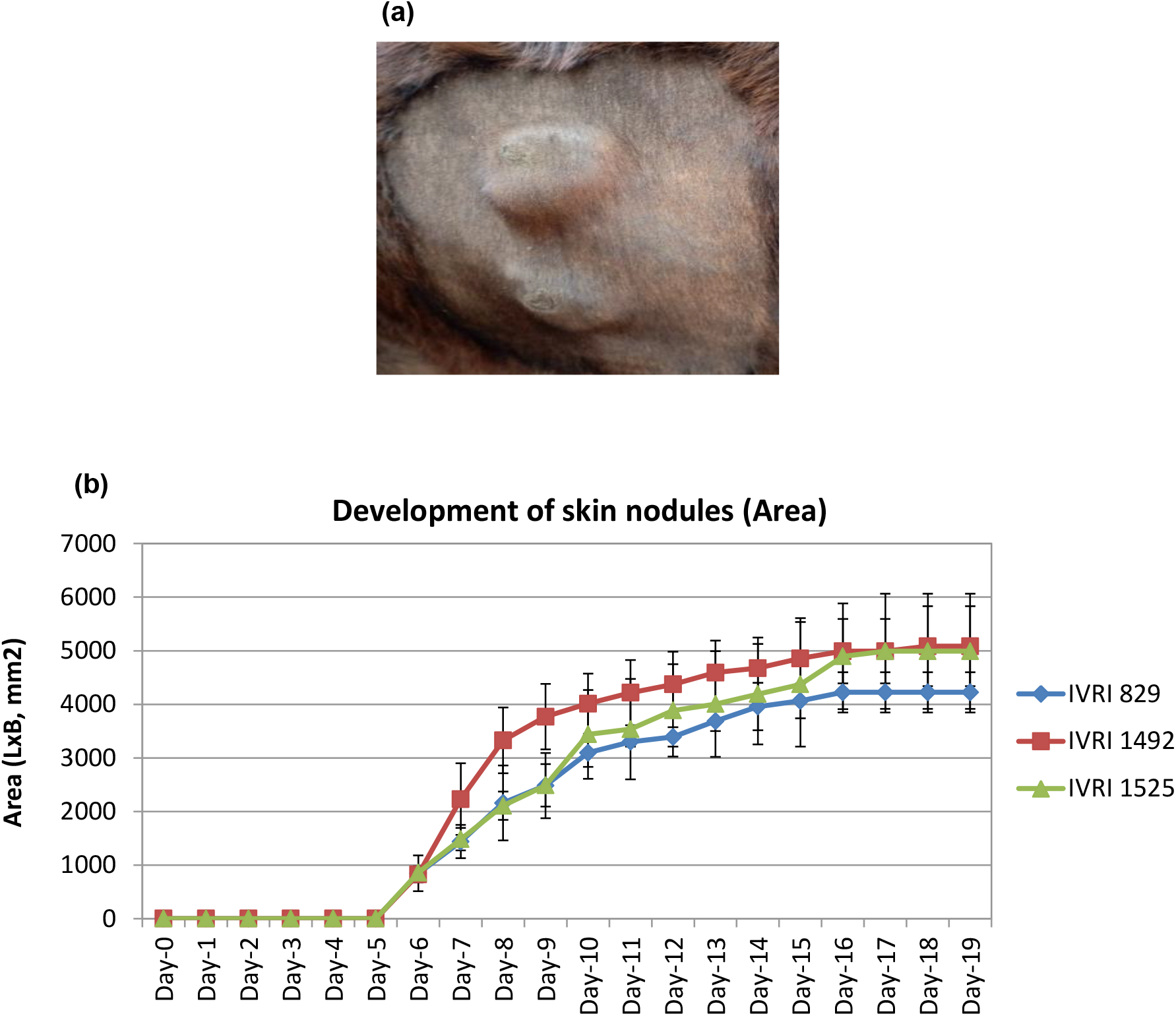
Development and progression of skin nodule following challenge with virulent LSDV. Animals were either unvaccinated or vaccinated with Lumpi-ProVac^Ind^. At day 30 pv, all the animals were challenged with virulent LSDV. Whereas vaccinated animals remained apparently healthy without showing any skin nodules, control (unvaccinated) animals developed the skin nodules following challenge **(a).** Primary swelling was seen at day 5-6 pc, which progressively increased in size from ~600 mm^2^ (day 6 pc) to 5000 mm^2^ (day 16 pc) before becoming stable at day 17 pc **(b).**

### Observations on animals vaccinated with Lumpi-ProVac^Ind^ in field

The LSD has become endemic in India. The outbreaks recorded have been very extensive with high morbidity and unusually high mortality. The field trials were initiated in clean (LSD free) herds during July-September 2022 and the animals were apparently healthy at the time of vaccination. Assuming that this is the first year of introduction of the disease (may be potentially free from maternal antibodies), we immunized all the animals including calves. Out of the 26940 animals across 102 vaccinated farms, there was no incidence of the disease till December 9, 2022 [except 14 animals (9 calves and 5 adults) across 5 farms wherein a mild disease was recorded] **(Table 2)**, despite the fact that severe disease with significant mortality was observed in the nearby unvaccinated farms.

### Virus excretion in milk and semen

We also evaluated the excretion of vaccine virus in milk of the vaccinated cows in the field. Paired milk sample, collected from day 3 to day 14 pv were found negative for LSDV genome by qRT-PCR **(Fig 5a** and **5b).** Likewise, samples (n=7) from bull semen collected on day 10 post-vaccination were also found negative for LSDV genome **(Fig 5c)**.

**Fig. 5.**
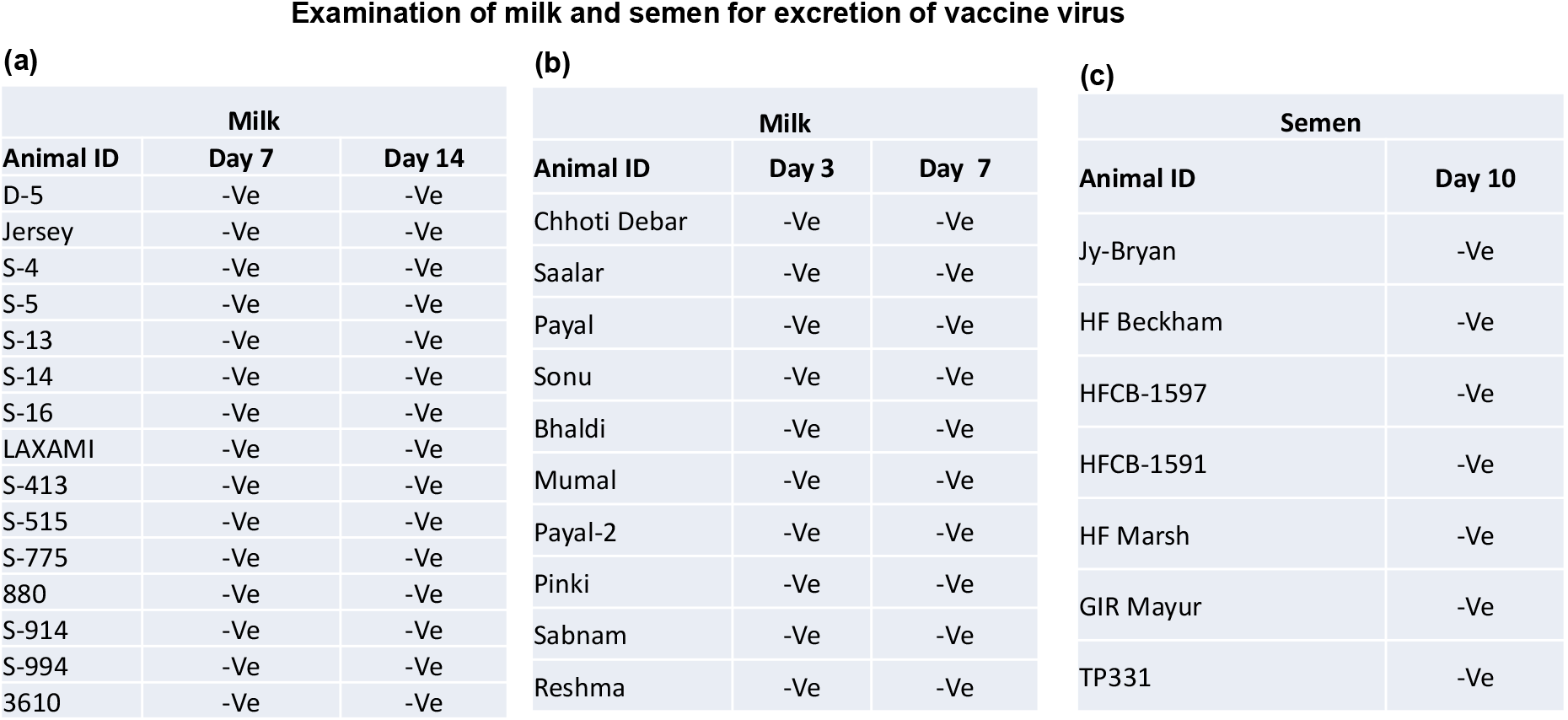
Evaluation of the excretion of vaccine virus in milk and semen in vaccinated animals under field conditions. Animals were vaccinated with Lumpi-ProVac^Ind^. Milk and semen samples were collected from the lactating cows and bulls respectively, and examined for the presence of LSDV genome. –Ve represents absence of LSDV genome.

### Exposure to virulent LSDV before development of complete immunity in endemic areas

A complete immunity usually develops 3-4 weeks following LSD vaccination^34, 35^. Since the disease was rampant and extensive outbreaks were being recorded in the surroundings farms/villages during the vaccine trials, vaccinated animals could have been exposed to the field/virulent virus. We observed the disease in 16 farms (in addition to 102 farms described above) during early times post-vaccination/during incubation period **(Table 4).** This was essentially due to insufficient immunity. In such cases, we were able to detect the field-but not vaccine strain of LSDV from the skin nodules **(Fig. 6)** which suggested the association of field virus as the cause of the disease. The overall morbidity and case fatality rate in these 16 farms was observed to be 7.1% and 2.71% respectively (**Table 4)**.

**Table 4:** Vaccinatioon during incubation period or early times post-infection.

| S. N. | Farm/Owner name | Total number of animals | Days after vaccination | Number of animals showed LSD symptoms | Total deaths |
| --- | --- | --- | --- | --- | --- |
| 1 | Dairy farm Rebari (Udaipur) | 47 | Day 3 | 4 | 0 |
| 2 | Gaurishankar Nagada | 95 | Day 4 | 2 | 0 |
| 3 | Rajesh Paliwal | 75 | Day 14 | 5 | 0 |
| 4 | Himmat Singh Naya Kheda | 40 | Day 10 | 15 | 0 |
| 5 | Laveshji | 40 | Day 10 | 3 | 0 |
| 6 | Ashokji Mali | 70 | Day 3 | 8 | 3 |
| 7 | Bhagwati lal Patodar | 41 | Day 10 | 5 | 0 |
| 8 | Viram Dev Singh | 40 | Day 4 | 1 | 0 |
| 9 | Dabok farm, Udaipur | 114 | Day 4 | 9 | 0 |
| 10 | Nutan Organic dairy farm, Jodhpur | 122 | Day 3 | 14 | 2 |
| 11 | Adrish Bhati Nohar, Raj | 63 | Day 6 | 6 | 0 |
| 12 | Narayan Singh/Prahald Ram Nagaur | 50 | Day 2 | 14 | 1 |
| 13 | Shri Shyam Gaushala, Talwana, Alwar | 381 | Day 7 | 11 | 0 |
| 14 | Himmat Singh, Dolat Singh Naya Kheda, Udaipur | 143 | Day 10 | 10 | 0 |
| 15 | Dwarkadhish Gaushala Talwara, Banswara | 200 | Day 3 | 8 | 0 |
| 16 | Anomol Gir Gaushala, Banswara | 150 | Day 2 | 4 | 0 |
| **Total number of animals** |  | **1671** |  | **119** | **6** |
| **Morbidity/Case fatality rate** |  |  |  | **Morbidity= 7.1%** | **CFR= 2.71%** |

**Fig. 6.**
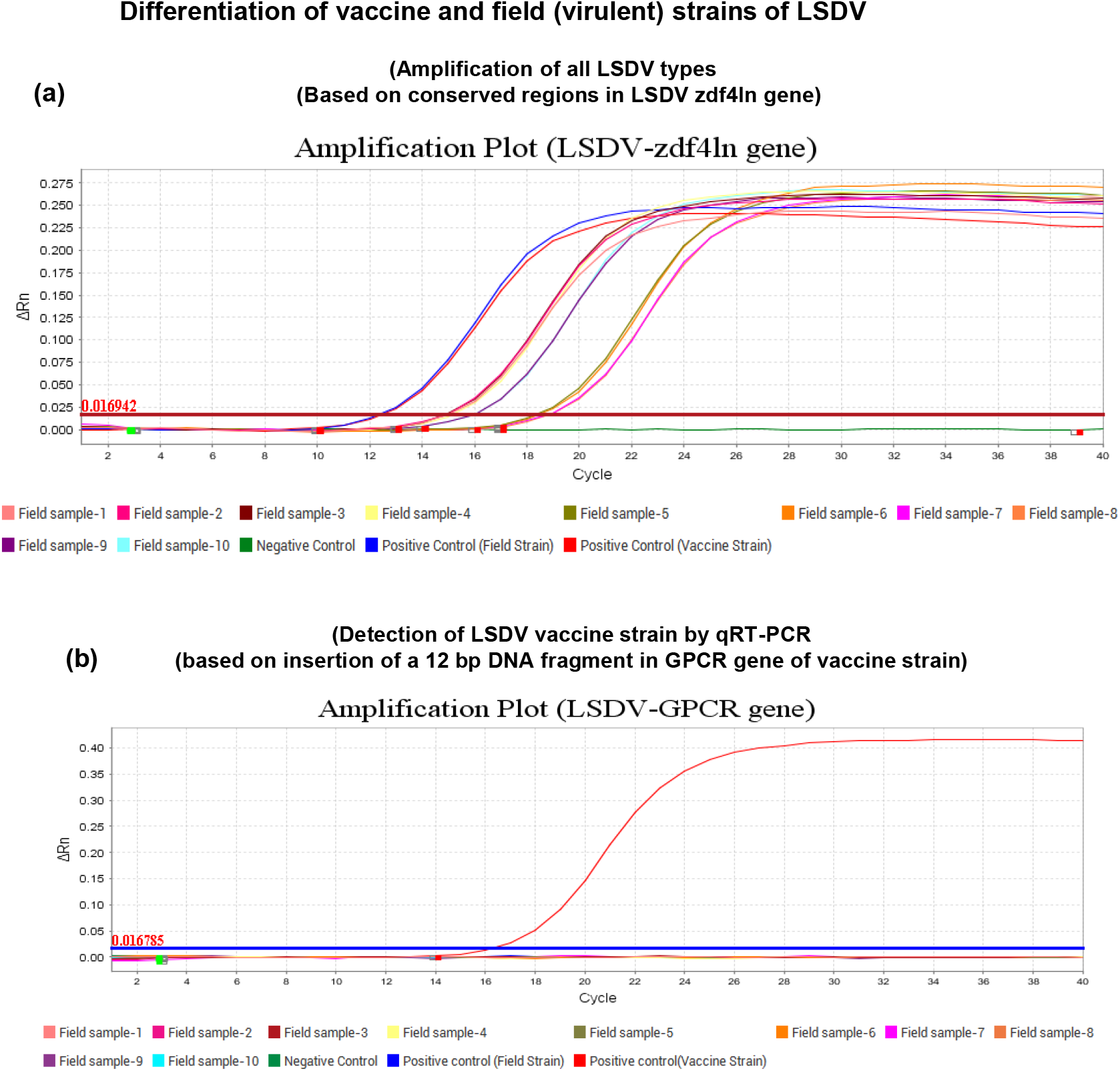
Differentiation of vaccine and field strains of LSDV. ***(a) Amplification of LSDV-specific gene (ORF44) segment***. Conserved regions of ORF44, present in all LSDV strains but not in other capripoxviruses (SPV/GPV) which was amplified by TaqMan real-time PCR, as described in the Materials and Method section. The amplification plots of the LSDV-Vaccine strain and LSDV-Field strains (n=10) are shown. ***(b) Amplification of Vaccine strain by TaqMan real-time PCR.*** A twelve bps insertion in LSDV GPCR gene, exclusively present in the vaccine strain but not in the field strain was exploited to differentiate the vaccine strains from the field strain(s). The amplification plot of the LSDV/Ranchi/P50 (vaccine strain) is shown. No amplification could be detected in the field strains (n=10).

### Vaccination after infection

Some of the animals that had developed the disease were also vaccinated in a limited number of animals on selected farms. The case fatality rate in such instances was recorded to be 0.09% **(Table 5).**

**Table 5:**
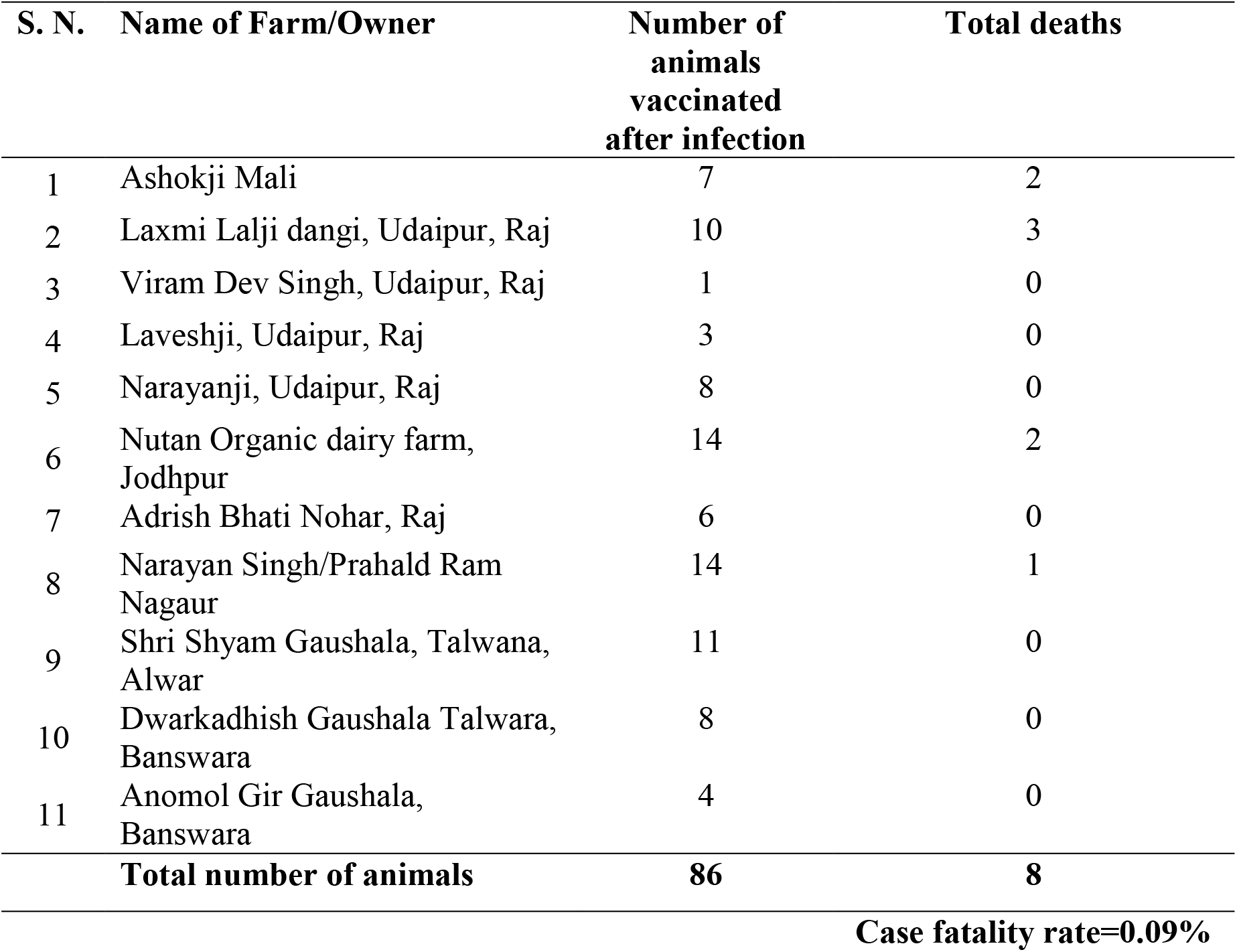
Vaccination after infection.

### Cross neutralization of LSDV strains

Sera derived from LSDV vaccinated (LSDV/2019/Ranchi/P50-vaccine strain)- or LSDV infected (LSDV/2019/Ranchi/P2-virulent strain) cattle, were subjected for their ability to neutralize different LSDV strains. These sera were able to equally neutralize various LSDV strains, irrespective of their history (2019 or 2022) or species of origin (camel or cattle) **(Table 6).**

**Table 6:** Neutralization of various LSDV strains by sera derived from LSDV/2019.

| **Virus** | **Sera derived from LSDV 2019** |  |  |  |  |  |
| --- | --- | --- | --- | --- | --- | --- |
|  | *Vaccinated sera* |  |  | *Infected sera* |  |  |
|  | **IVRI-1489** | **IVRI-1524** | **IVRI-1533** | **IVRI-1525** | **IVRI-829** | **IVRI-1492** |
| LSDV/Cattle/India/2019/Ranchi | 64 | 32 | 32 | 32 | 32 | 32 |
| LSDV/Cattle/India/2021/Banswara | 32 | 64 | 32 | 32 | 16 | 32 |
| LSDV/Cattle/India/2022/Jalore | 64 | 32 | 32 | 32 | 32 | 32 |
| LSDV/**Camel**/India/2022/Bikaner | 64 | 32 | 64 | 16 | 32 | 32 |

Values represents neutralization titre

## Discussion

India is currently facing a deadly epidemic of LSD. Over 125,000 cattle have died and around 20 million cattle have been infected^1^. Capripoxviruses are genetically quite similar and antigenically indistinguishable; therefore all fall under a single serotype^36, 37^. Due to potential safety issues, the affected countries usually tend to authorize only those *Capripoxvirus* vaccines against which infecting viruses are prevalent^4^. Due to the unavailability of a homologous LSD vaccine, the policy makers in India decided to use the heterologous (GPV-based) vaccine to induce immunoprophylaxis against LSDV in cattle^18^ as an emergency measure. However, heterologous vaccines provide only partial protection against LSDV in cattle and are not as efficacious as homologous LSDV-based vaccine, therefore necessitating a homologous LSD vaccine for vaccination in cattle^13, 21–25, 38, 39^. The discrepancies and inconsistency of results obtained from many heterologous vaccines against LSD prompted us to develop a homologous vaccine.

The virus was attenuated by continuous passages in Vero cells. As compared to the field/virulent strain (LSDV/Ranchi/P0), the whole genome sequencing of the vaccine strain (LSDV/Ranchi/P50) revealed several mutations. The major mutations were observed in DNA-dependent RNA polymerase, Kelch-like protein, EEV membrane phospho-glycoprotein and CD47-like putative membrane protein, most of which have been reported in other *Capripoxvirus* vaccine strains as well^33^. A twelve bps insertion in the GPCR gene, which is considered as a signature mutation in vaccine strains^30^ was observed even at P10 level and was maintained in subsequent passages in our study. In addition, Ranchi/P50 had an 801 bp deletion in its inverted terminal repeat (ITR) region which has not been reported in any other vaccine/field strains. This major deletion could also be consistently detected at passage levels, P40, P50, P60 and P70.

As compared to the classical Neethling LSDV strains which require over 100 passages for attenuation^40^, KSGP strains require relatively low number of passages (~30) for attenuation^34^. Indian LSDV strains, including LSDV/India/2019/Ranchi which was used to prepare the vaccine is a NI-2490/Kenya/KSGP-like field strains (somewhere in between Neethling and KSGP LSDV strains)^9, 41^. This, together with the similarities of LSDV/Ranchi/P50 with other *Capripoxvirus* vaccine strains tempted us to speculate about the potential attenuation of the P50 virus, and, therefore paving the way to conduct vaccine safety and efficacy studies in experimental calves.

The vaccine was well tolerated in calves with no local or systemic reaction or any other adverse effect on feed uptake, behaviour and general health status being observed. Also, the shedding of the vaccine virus was not detected in ocular, nasal and faecal swab from day 1 to day 30 pv. Further, viremia (virus isolation from blood) could not be detected in any of the vaccinated animals, except that few of the animals had low copy number of viral genome at day 3 post-vaccination. Besides, various blood biochemical parameters were also normal. Taken together, experimental inoculation of the vaccine in cattle was found to be safe.

Upon challenge infection, all the unvaccinated (naïve) control animals developed localized skin nodules (onset at day 5-6 pc), besides developing fever and viremia between day 7 and day 10 pc which lasted for 1-2 days. The unvaccinated control animals were also anorectic and depressed at day 7-8 pc. These data were comparable to previous studies on the efficacy of homologous LSD vaccines^37, 42–46^. On the other hand, vaccinated-challenged animals completely resisted the development of skin nodules, fever and viremia which strongly suggests that the LSDV/Ranchi/P50-based vaccine provides protective (100%) efficacy against virulent LSDV.

Homologous live-attenuated vaccines are considered to be most immunogenic and efficacious. Neethling LSDV strain from South Africa has primarily been used as a homologous LSDV strain in most part of the world^47^. Currently, at least eleven LSD vaccines have been described and are being used in different parts of the world to induce immunoprophylaxis against LSD in cattle^34^. Most of the homologous LSDV vaccines (Lumpyvax™, Bovivax-LSD^™^, LumpyShield-N^™^ and MEVAC LSD) contain the well-known South African Neethling strain, despite their different passage/attenuation history^34^. Neethling strain-based vaccine was recently used to contain the disease in Balkan countries with an effectiveness of >80%^48^. The use of Kenyan sheep and goat pox (KSGP) virus strains O-240 and O-180 (strains are of ovine origin, isolated during the same outbreak but at different points in time) which were later confirmed as LSDV strains^17, 49, 50^ has been limited to Egypt^51^, Ethiopia^52^ and Israel^5^. Besides, heterologous vaccines based on Yugoslavian RM65 sheep pox (SPP), Romanian SPP and Gorgan goat pox (GPT) strains have also been used in cattle against LSD^34^. Homologous live-attenuated LSD vaccines described above are known to cause adverse effects in cattle. This includes swelling at the site of vaccination or rarely generalized small-size skin nodules and a temporal reduction in the milk yield which is often referred to as “Neethling disease” or “Neethling response“^46, 53, 54^. In our study, none of the 10 vaccinated animals including 2 animals that received 10-times of field dose (10^4.5^ TCID_50_) developed any local or systemic reaction (nodule) following vaccination (experimental trial). In field conditions, only 0.018% of the animals (5 out of the 26940 animals vaccinated) showed local site reaction (appeared at day 3 pv and subsided within 2 days) whereas generalized skin nodules were not reported in any of the vaccinated animals. In the previous studies under field conditions, the percentage of vaccinated animals exhibiting LSD-like nodules after vaccination was shown to vary from .09% to 12%^26, 34, 54, 55^. Besides, vaccine dose and immune status of the vaccinated animals, the nature of the vaccine strain used could be major factors for the Neethling response^34^. While harbouring most of the mutations reported in other *Capripoxvirus* vaccine strains, LSDV/Ranchi (close to Kenyan LSDV/KSGP strain) has a unique deletion of 801 nucleotides in its ITR region. This unique deletion mutation might be associated with extremely low levels of Neethling response in LSDV/Ranchi/P50-based vaccine, although further experimental proof will be essential.

A fever response was usually observed in animals following vaccination with Neethling strain^37, 42–46^. In our study, rise in temperature following vaccination was recorded in a total of four animals viz; at day 12 to day 16 pv (four animals) and at day 1 to day 5 pv (two animals). We could not ascertain whether this rise in temperature is specific in response to vaccine or else due to other biotic or abiotic stress factors, because there was no specific pattern (onset and duration) of fever. Nevertheless, like in other existing LSDV and other viral vaccines, fever for a few days in some of the vaccine recipients is a common phenomenon^34^.

In natural infection, the infected animals may excrete virus in the milk^56^ and semen^57^. Although there is no confirmed report on excretion of LSDV vaccine virus in the semen ^57^, a small percentage of the vaccinated cows may excrete virus in the milk^56, 58^. In our study, we neither observed excretion of the LSDV/Ranchi/P50 (vaccine virus) in milk, nor in semen. This suggests that the milk from vaccinated animals (Lumpi-ProVac^Ind^) does not present any harm to public health. Likewise, there is no risk of vertical transmission of vaccine virus via bull semen.

The seroconversion in our study was seen as early as day 18 pv. All the 10 vaccinated animals, except one, developed anti-LSDV antibodies (seroconversion rate 90%) on the day 30 pv. In the field studies, seroconversion rate was 85.18% (n=648) at day 30 pv.

Seroconversion rate following LSD vaccination is highly variable (40-80%)^59–62^. Since LSD has become endemic in India and extensive outbreaks were being recorded at the time of the vaccine trials, it is likely that the high seroconversion rate is presumably due to exposure to natural infection, post-vaccination. This essentially necessitates the development of a test system to differentiate among the antibodies generated due to infection or vaccination.

Although the under trial animals were tested negative for antibodies against capripoxviruses before start of the safety and efficacy trials, but one of the vaccinated animal (IVRI 1462) and two (IVRI-835 and IVRI-1499) of the five control animals had anti-LSDV antibodies at the time of vaccination (day 0 pv), and challenge (day 0 pc) infection respectively. Unvaccinated control animals with pre-existing antibodies (unknown history of LSDV exposure) at the time of challenge infection were completely protected upon virulent challenge. This suggests that they were essentially exposed to LSDV in between screening (for seronegativity) and the time of challenge. Interestingly, these animals also exhibited an IFY-ϒ response in serum. Whether increased IFY-ϒ in these animals was due to LSDV exposure or due to cryptic infections^63^ could not be ascertained.

Lymphoproliferation and levels of IFN-ϒ were estimated as a measure of cell-mediated immunity (CMI). Significant cell proliferation was observed in 60% of the vaccinated calves at day 30 pv. In agreement with the previous studies^59, 60, 64^, the peak response of IFN-ϒ was observed at day 3 post LSDV exposure (irrespective of vaccination/infection), the highest being in vaccinated-challenged (100%) than in unvaccinated-challenged (66%) or merely vaccinated (50%) animals. However, the precise role of CMI in protection against LSDV is poorly understood and needs further investigations.

We did not observe any correlation between the antibody titres and cell-mediated immune response. Animals that developed the highest antibody titres did not show significant cell-mediated immune response. Likewise, one animal that did not develop antibodies (at day 30 pv), showed cell proliferation response. However, irrespective of their immune status, all the vaccinated animals resisted the challenge with the virulent LSDV. With the reasons unknown, protection from virulent LSDV in the absence of detectable anti-LSDV neutralizing antibodies has also been reported by other workers^21, 46, 57, 65^. While antibody response following vaccination of cattle with LSDV is variable (40-80%), the antibodies may be undetectable following administration of SPV vaccines ^20, 66^. This may partly be attributed to the nature of some cattle breeds or the type of vaccine used^67, 68^. In a study by Aboul Soud *et al.*, no serological response was induced in cattle vaccinated with Romania strain while a trivalent Capripox vaccine (made up of SP Romania, GTPV Held and KSGP 0180) induced antibodies in 66% of vaccinated animals^66^. The mechanism underlying the protection of vaccinated animals do not developing antibodies and/or CMI (Lymphoproliferation/IFN response) at the time of challenge needs further investigation.

Although none of the live-attenuated capripoxvirus vaccines have been claimed to provide total protection, however, the spread of LSD has been successfully controlled using Neethling vaccines at high levels of vaccine coverage^4^. LSD vaccines are widely used in Africa, although vaccine breakdown and reinfection of vaccinated animals have been reported^69^. The heterogeneity of viruses used in vaccine, use of over attenuated virus strain and inappropriate production process (quality) of the vaccine may lead to failure in the generation of protective immunity^21, 22, 34, 42^. However, with a vaccine effectiveness of up to 97%, Neethling vaccine was recently shown to be highly effective, in controlling the LSD epidemic in six Balkan countries ^70^. Although our analysis include the observations for maximum of 4 months post-vaccination, the vaccine efficacy was determined to be 99.94% [out of 26940 animals, only 14 animals (0.05%) developed the disease after completion of 21 days of vaccination]. The numbers of calves included for vaccination in our study were <5%, however, majority of the total cases of vaccine ineffectiveness in our study were observed in calves (52.52%). The poor efficacy in calves might be attributed to interference of vaccination by maternal antibodies^26, 71^. However, this also needs further investigation.

The field trials were initiated in clean disease free herds. However, since the disease was widely prevalent, some of the animals exhibited symptoms of LSD early post-vaccination, which was presumably due to encountering the natural infection and was confirmed by differentiating the infecting (field) virus with vaccine virus by PCR. Since full protective immunity is believed to develop only after 3-4 weeks post-vaccination^34, 35^, animals under incubation period or before completion of 3 weeks of vaccination may develop the disease as expected. All such cases were observed in Rajasthan which was the most badly affected state. In such instances, the morbidity and case fatality rate was 7.1% and 2.71% respectively **(Table 4),** which was significantly low as compared to the overall morbidity and case fatality rate (11.28% and 4.8% respectively) in the state^72^. Therefore, our findings appear to suggest practice of vaccinating animals with LSD vaccine in outbreak affected areas, irrespective of the history of the animals in close contact with the infected animals. However, there are conflicting reports on outcome of the disease in animals that succumb to natural infection at early times post-vaccination. A study by Ayelet suggests high morbidity in local breed but no significant effect in cross breeds ^52^. Study by Abutrabush suggested highest morbidity and mortality in the order of nonvaccinated farms>vaccinated farms after infection>vaccinated farms^22^. It is speculated that while the virulent virus is preparing for tissue damage during disease progression, some immunity elicited by the vaccine virus could neutralise the virulent virus, thus slowing the disease process. However, this aspect of pathogenesis and protection mechanism in infected and vaccinated animals needs experimental proof and in-depth investigation.

The average case fatality rate in the animals that were vaccinated after appearance of the disease was low (0.09%) as compared to the overall case fatality rate (4.8%) in the state, which appears to suggest that in most instances it has a beneficial effect (reducing disease severity). However, the case fatality rate, which varied from 0-30%, also appears to suggest that in certain instances, vaccination of infected animals may have an adverse impact on disease severity. The beneficial or adverse impact may depend on the time of vaccinating infected animals, besides coinfection of other microbial agents^63^. This needs further investigation.

A perfect cross protection has been established among various capripoxviruses in general, and among the LSDV strains in particular ^1^. However, due to unusual high mortality involved in recent LSDV outbreak in Indian subcontinent, the concerns were raised about the protective efficacy of the vaccine. The sera derived from LSDV/2019 were able to equally neutralize LSDV/2019, LSDV/2021 and LSDV/2022. This, together with high efficacy of Lumpi-ProVac^Ind^ in the field (against LSDV/2022) suggests that the vaccine can be conveniently employed for control and eradication of LSD.

## Conclusion

The safety profile of Lumpi-ProVac^Ind^ is very high (minimal or no Neethling response) as compared to the other existing live-attenuated vaccines. It was found to be highly efficacious against LSD and could prove to be a better option for the control and eradication of LSD in India as well as other affected countries.

## Data availability

All the data are available within this manuscript. LSDV/Cattle/India/2021/Banswara, LSDV/Cattle/India/2022/Jalore and LSDV/Camel/India/2022/Bikaner have been deposited at the National repository (NCVTC, Hisar India) with Accession Numbers of VTCCAVA 321, VTCCAVA 370 and VTCCAVA 371 respectively. The complete nucleotide sequences of LSDV/India/2019/Ranchi/P0 and LSDV/India/2019/Ranchi/P50 are available with GenBank Accession Numbers of MW883897.1 and OK422494.1, respectively.

## Disclosure statement

The authors claim no conflict of interest exits in the submission of this manuscript. The manuscript has been approved by all authors for publication. The work is original research that has not been published previously, and not under consideration for publication elsewhere, in whole or in part.

## Author contributions

N.Ku., B.N.T., S.B. and T.D. conceived and designed the research. N.Ku., S.B., Rm.Ku., N.Kh., B.G., Y.C., S.K., R.K.D. and A.P. collected samples performed laboratory experiments. N.Ku., S.B., Rm.Ku., N.Kh., A.V., A.K., L.S., S.N., S.G., A.G., As.K., J.M., K.P.S., Y.P., T.D. and A.K.M. participated in experimental trial. N.Ku., S.B., Rm.Ku., N.Kh., A.V., L.S., D.K.S., V.M., B.B., R.T., V.V., S.C., V.Y., Ab.B., R.Ka., A.B., A.A., R.W.Y., A.K., S.K., H.A.T., M.K.G., Rj.Ku. and Y.P. performed field trials. N.Ku., B.N.T., Rm.Ku. and S.B. analysed the data. N.Ku. wrote the primary draft of manuscript. All authors reviewed the manuscript.

## Funding

This work was supported by Indian Council of Agricultural Research, New Delhi [grant number IXX11882] and Science and Engineering Research Board, Department of Science and Technology, Government of India [grant number CRG/2018/004747].

## Supplementary Materials

**Table S1:** Body temperature of vaccinated calves (Experimental trial). **Table S2:** Body temperature of vaccinated cattle (Field trial). **Table S3:** Body temperature of vaccinated buffaloes (Field trial). **Table S4:** Body temperature of vaccinated and unvaccinated calves following challenge infection (Experimental trial).

## Notes

### Competing Interest Statement

The authors have declared no competing interest.

## References

1. Kumar N, Tripathi BN. A serious skin virus epidemic sweeping through the Indian subcontinent is a threat to the livelihood of farmers. Virulence 2022; 13:1943–4.

2. CABI. Lumpy skin Disease. https://www.cabiorg/isc/datasheet/76780 2019.

3. Ras M. Pseudo-Urticaria of Cattle. Government of Northern Rhodesia: Department of Animal Health 1931:20–1.

4. Tuppurainen ES, Oura CA. Review: lumpy skin disease: an emerging threat to Europe, the Middle East and Asia. Transbound Emerg Dis 2012; 59:40–8.

5. Yeruham I, Nir O, Braverman Y, Davidson M, Grinstein H, Haymovitch M, et al. Spread of lumpy skin disease in Israeli dairy herds. Vet Rec 1995; 137:91–3.

6. Sudhakar SB, Mishra N, Kalaiyarasu S, Jhade SK, Hemadri D, Sood R, et al. Lumpy skin disease (LSD) outbreaks in cattle in Odisha state, India in August 2019: Epidemiological features and molecular studies. Transbound Emerg Dis 2020.

7. Tran HTT, Truong AD, Dang AK, Ly DV, Nguyen CT, Chu NT, et al. Lumpy skin disease outbreaks in vietnam, 2020. Transbound Emerg Dis 2021; 68:977–80.

8. Roche X, Rozstalnyy A, TagoPacheco D, Pittiglio C, Kamata A, Beltran Alcrudo D, et al. Introduction and spread of lumpy skin disease in South, East and Southeast Asia: Qualitative risk assessment and management. Rome, FAO.: FAO animal production and health, 2020.

9. Kumar N, Chander Y, Kumar R, Khandelwal N, Riyesh T, Chaudhary K, et al. Isolation and characterization of lumpy skin disease virus from cattle in India. PLoS One 2021; 16:e0241022.

10. Casal J, Allepuz A, Miteva A, Pite L, Tabakovsky B, Terzievski D, et al. Economic cost of lumpy skin disease outbreaks in three Balkan countries: Albania, Bulgaria and the Former Yugoslav Republic of Macedonia (2016-2017). Transbound Emerg Dis 2018; 65:1680–8.

11. Shagun. Lump Skin Disease has created a livehood crisis for India’s small dairy farmers. DownToEarth, September 30 2022:(https://www.downtoearth.org.in/news/economy/ground-report-lumpy-skin-disease-has-created-a-livelihood-crisis-for-india-s-small-dairy-farmers-85245).

12. Lojkic I, Simic I, Kresic N, Bedekovic T. Complete Genome Sequence of a Lumpy Skin Disease Virus Strain Isolated from the Skin of a Vaccinated Animal. Genome Announc 2018; 6.

13. Abutarbush SM, Tuppurainen ESM. Serological and clinical evaluation of the Yugoslavian RM65 sheep pox strain vaccine use in cattle against lumpy skin disease. Transbound Emerg Dis 2018; 65:1657–63.

14. Tuppurainen ESM, Venter EH, Shisler JL, Gari G, Mekonnen GA, Juleff N, et al. Review: Capripoxvirus Diseases: Current Status and Opportunities for Control. Transbound Emerg Dis 2017; 64:729–45.

15. Teffera M, Babiuk S. Potential of Using Capripoxvirus Vectored Vaccines Against Arboviruses in Sheep, Goats, and Cattle. Front Vet Sci 2019; 6:450.

16. Kitching RP. Vaccines for lumpy skin disease, sheep pox and goat pox. Dev Biol (Basel) 2003; 114:161–7.

17. Tuppurainen ES, Pearson CR, Bachanek-Bankowska K, Knowles NJ, Amareen S, Frost L, et al. Characterization of sheep pox virus vaccine for cattle against lumpy skin disease virus. Antiviral Res 2014; 109:1–6.

18. AnnualReport. Guidelines and advisory on lumpy skin disease virus. Annual Report of Department of Animal Husbandry, Dairying and Fisheries, Ministry of Animal Husbandry and Dairying, Government of India 2022:pp 89.

19. Gaber A, Rouby S, Elsaied A, El-Sherif A. Assessment of heterologous lumpy skin disease vaccine-induced immunity in pregnant cattle vaccinated at different times of gestation period and their influence on maternally derived antibodies. Vet Immunol Immunopathol 2022; 244:110380.

20. Hamdi J, Boumart Z, Daouam S, El Arkam A, Bamouh Z, Jazouli M, et al. Development and Evaluation of an Inactivated Lumpy Skin Disease Vaccine for Cattle. Vet Microbiol 2020; 245:108689.

21. Brenner J, Bellaiche M, Gross E, Elad D, Oved Z, Haimovitz M, et al. Appearance of skin lesions in cattle populations vaccinated against lumpy skin disease: statutory challenge. Vaccine 2009; 27:1500–3.

22. Abutarbush SM. Efficacy of vaccination against lumpy skin disease in Jordanian cattle. Vet Rec 2014; 175:302.

23. Sevik M, Dogan M. Epidemiological and Molecular Studies on Lumpy Skin Disease Outbreaks in Turkey during 2014-2015. Transbound Emerg Dis 2017; 64:1268–79.

24. Rao TV, Bandyopadhyay SK. A comprehensive review of goat pox and sheep pox and their diagnosis. Anim Health Res Rev 2000; 1:127–36.

25. Abd-Elfatah EB, El-Mekkawi, M. F. & Aboul-Soud, E. A. Capripoxviruses of small ruminants: control and evaluating the future update efficacy of a current vaccine in Egypt. Advances in Environmental Biology 2018; 12:11–6.

26. Ben-Gera J, Klement E, Khinich E, Stram Y, Shpigel NY. Comparison of the efficacy of Neethling lumpy skin disease virus and x10RM65 sheep-pox live attenuated vaccines for the prevention of lumpy skin disease - The results of a randomized controlled field study. Vaccine 2015; 33:4837–42.

27. Kumar N, Wadhwa A, Chaubey KK, Singh SV, Gupta S, Sharma S, et al. Isolation and phylogenetic analysis of an orf virus from sheep in Makhdoom, India. Virus Genes 2014; 48:312–9.

28. Khandelwal N, Chander Y, Rawat KD, Riyesh T, Nishanth C, Sharma S, et al. Emetine inhibits replication of RNA and DNA viruses without generating drug-resistant virus variants. Antiviral Res 2017; 144:196–204.

29. Alexander S, Olga B, Svetlana K, Valeriy Z, Yana P, Pavel P, et al. A real-time PCR screening assay for the universal detection of lumpy skin disease virus DNA. BMC Res Notes 2019; 12:371.

30. Agianniotaki EI, Chaintoutis SC, Haegeman A, Tasioudi KE, De Leeuw I, Katsoulos PD, et al. Development and validation of a TaqMan probe-based real-time PCR method for the differentiation of wild type lumpy skin disease virus from vaccine virus strains. J Virol Methods 2017; 249:48–57.

31. WOAH. Manual of Diagnostic Tests and Vaccines for Terrestrial Animals 2021. Lumpy Skin Disease, Chapter 3.4.12. 2021; Retrieved from https://www.woah.org/fileadmin/Home/eng/Health_standards/tahm/3.04.12_LSD.pdf (Accessed on December 8, 2022).

32. Sanchez-Sampedro L, Perdiguero B, Mejias-Perez E, Garcia-Arriaza J, Di Pilato M, Esteban M. The evolution of poxvirus vaccines. Viruses 2015; 7:1726–803.

33. Biswas S, Noyce RS, Babiuk LA, Lung O, Bulach DM, Bowden TR, et al. Extended sequencing of vaccine and wild-type capripoxvirus isolates provides insights into genes modulating virulence and host range. Transbound Emerg Dis 2020; 67:80–97.

34. Tuppurainen E, Dietze K, Wolff J, Bergmann H, Beltran-Alcrudo D, Fahrion A, et al. Review: Vaccines and Vaccination against Lumpy Skin Disease. Vaccines (Basel) 2021; 9.

35. Woods JA. Lumpy skin disease--a review. Trop Anim Health Prod 1988; 20:11–7.

36. Hamdi J, Munyanduki H, Omari Tadlaoui K, El Harrak M, Fassi Fihri O. Capripoxvirus Infections in Ruminants: A Review. Microorganisms 2021; 9.

37. Babiuk S, Bowden TR, Parkyn G, Dalman B, Manning L, Neufeld J, et al. Quantification of lumpy skin disease virus following experimental infection in cattle. Transbound Emerg Dis 2008; 55:299–307.

38. Hamdi J, Bamouh Z, Jazouli M, Boumart Z, Tadlaoui KO, Fihri OF, et al. Experimental evaluation of the cross-protection between Sheeppox and bovine Lumpy skin vaccines. Sci Rep 2020; 10:8888.

39. Mikhael C, Ibrahim M, Saad M. Efficacy of Alternative Vaccination with Attenuated Sheep Pox and Inactivated Lumpy Skin Disease Vaccines against Lumpy Skin Disease. Suez Canal Veterinary Medical Journal SCVMJ 2016; 21:125–42.

40. Chervyakova OV SK, Nissanova RK, Orynbayev MB. Lumpy Skin Disease Virus: Approaches to attenuation for vaccine development. Eurasian Journal of Applied Biotechnology 2019; 1:3–12.

41. Sudhakar SB, Mishra N, Kalaiyarasu S, Jhade SK, Hemadri D, Sood R, et al. Lumpy skin disease (LSD) outbreaks in cattle in Odisha state, India in August 2019: Epidemiological features and molecular studies. Transbound Emerg Dis 2020; 67:2408–22.

42. Gari G, Abie G, Gizaw D, Wubete A, Kidane M, Asgedom H, et al. Evaluation of the safety, immunogenicity and efficacy of three capripoxvirus vaccine strains against lumpy skin disease virus. Vaccine 2015; 33:3256–61.

43. Prozesky L, Barnard BJ. A study of the pathology of lumpy skin disease in cattle. Onderstepoort J Vet Res 1982; 49:167–75.

44. Moller J, Moritz T, Schlottau K, Krstevski K, Hoffmann D, Beer M, et al. Experimental lumpy skin disease virus infection of cattle: comparison of a field strain and a vaccine strain. Arch Virol 2019; 164:2931–41.

45. Sanz-Bernardo B, Haga IR, Wijesiriwardana N, Hawes PC, Simpson J, Morrison LR, et al. Lumpy Skin Disease Is Characterized by Severe Multifocal Dermatitis With Necrotizing Fibrinoid Vasculitis Following Experimental Infection. Vet Pathol 2020; 57:388–96.

46. Haegeman A, De Leeuw I, Mostin L, Campe WV, Aerts L, Venter E, et al. Comparative Evaluation of Lumpy Skin Disease Virus-Based Live Attenuated Vaccines. Vaccines (Basel) 2021; 9.

47. Davies FG. Lumpy skin disease of cattle: A growing problem in Africa and the Near East. World Animal Review 1991; 68:37–42.

48. Klement E, Broglia A, Antoniou S-E, Tsiamadis V, Plevraki E, Petrović T, et al. Neethling vaccine proved highly effective in controlling lumpy skin disease epidemics in the Balkans. Preventive Veterinary Medicine 2020; 181:104595.

49. Lamien CE, Le Goff C, Silber R, Wallace DB, Gulyaz V, Tuppurainen E, et al. Use of the Capripoxvirus homologue of Vaccinia virus 30 kDa RNA polymerase subunit (RPO30) gene as a novel diagnostic and genotyping target: development of a classical PCR method to differentiate Goat poxvirus from Sheep poxvirus. Vet Microbiol 2011; 149:30–9.

50. Tulman ER, Afonso CL, Lu Z, Zsak L, Sur JH, Sandybaev NT, et al. The genomes of sheeppox and goatpox viruses. J Virol 2002; 76:6054–61.

51. Salib F.A. OAH. Incidence of lumpy skin disease among Egyptian cattle in Giza Governorate, Egypt. Veterinary World 2011 4:162–7.

52. Ayelet G, Abate Y, Sisay T, Nigussie H, Gelaye E, Jemberie S, et al. Lumpy skin disease: preliminary vaccine efficacy assessment and overview on outbreak impact in dairy cattle at Debre Zeit, central Ethiopia. Antiviral Res 2013; 98:261–5.

53. Tuppurainen ESM, Antoniou SE, Tsiamadis E, Topkaridou M, Labus T, Debeljak Z, et al. Field observations and experiences gained from the implementation of control measures against lumpy skin disease in South-East Europe between 2015 and 2017. Prev Vet Med 2020; 181:104600.

54. Katsoulos PD, Chaintoutis SC, Dovas CI, Polizopoulou ZS, Brellou GD, Agianniotaki EI, et al. Investigation on the incidence of adverse reactions, viraemia and haematological changes following field immunization of cattle using a live attenuated vaccine against lumpy skin disease. Transbound Emerg Dis 2018; 65:174–85.

55. Abutarbush SM, Hananeh WM, Ramadan W, Al Sheyab OM, Alnajjar AR, Al Zoubi IG, et al. Adverse Reactions to Field Vaccination Against Lumpy Skin Disease in Jordan. Transbound Emerg Dis 2016; 63:e213–9.

56. Bedekovic T, Simic I, Kresic N, Lojkic I. Detection of lumpy skin disease virus in skin lesions, blood, nasal swabs and milk following preventive vaccination. Transbound Emerg Dis 2018; 65:491–6.

57. Osuagwuh UI, Bagla V, Venter EH, Annandale CH, Irons PC. Absence of lumpy skin disease virus in semen of vaccinated bulls following vaccination and subsequent experimental infection. Vaccine 2007; 25:2238–43.

58. Namazi F, Khodakaram Tafti A. Lumpy skin disease, an emerging transboundary viral disease: A review. Vet Med Sci 2021; 7:888–96.

59. Norian R, Afzal Ahangaran N, Azadmehr A. Evaluation of humoral and cell-mediated immunity of two capripoxvirus vaccine strains against lumpy skin disease virus. Iran J Virol 2016; 10:1–11.

60. Varshovi HR, Norian R, Azadmehr A, Afzal Ahangaran N. Immune response characteristics of Capri pox virus vaccines following emergency vaccination of cattle against lumpy skin disease virus. Iranian Journal of Veterinary Science and Technology 2017; 9:33–40.

61. Bhanuprakash V, Indrani BK, Hegde R, Kumar MM, Moorthy AR. A classical live attenuated vaccine for sheep pox. Trop Anim Health Prod 2004; 36:307–20.

62. Boumart Z, Daouam S, Belkourati I, Rafi L, Tuppurainen E, Tadlaoui KO, et al. Comparative innocuity and efficacy of live and inactivated sheeppox vaccines. BMC Vet Res 2016; 12:133.

63. Kumar N, Sharma S, Barua S, Tripathi BN, Rouse BT. Virological and Immunological Outcomes of Coinfections. Clin Microbiol Rev 2018; 31.

64. Khafagy HA, Saad MA, Abdelwahab MG, Mustafa AM. Preparation and field evaluation of live attenuated sheep pox vaccine for protection of calves against lumpy skin disease. Benha Veterinary Medical Journal 2016; 31:1–7.

65. Abutarbush SM, Ababneh MM, Al Zoubi IG, Al Sheyab OM, Al Zoubi MG, Alekish MO, et al. Lumpy Skin Disease in Jordan: Disease Emergence, Clinical Signs, Complications and Preliminary-associated Economic Losses. Transbound Emerg Dis 2015; 62:549–54.

66. Kafafy M, El Soally S, Aboul-Soud E, Zaghloul M, Mikhael C. Preparation of trivalent vaccine against lumpy skin disease using different capripox viral strain. Veterinary Medical Journal (Giza) 2018; 64:23–38.

67. Abdallah FM, El Damaty HM, Kotb GF. Sporadic cases of lumpy skin disease among cattle in Sharkia province, Egypt: Genetic characterization of lumpy skin disease virus isolates and pathological findings. Vet World 2018; 11:1150–8.

68. Bamouh Z, Hamdi J, Fellahi S, Khayi S, Jazouli M, Tadlaoui KO, et al. Investigation of Post Vaccination Reactions of Two Live Attenuated Vaccines against Lumpy Skin Disease of Cattle. Vaccines 2021; 9:621.

69. Kara PD, Afonso CL, Wallace DB, Kutish GF, Abolnik C, Lu Z, et al. Comparative sequence analysis of the South African vaccine strain and two virulent field isolates of Lumpy skin disease virus. Arch Virol 2003; 148:1335–56.

70. Klement E, Broglia A, Antoniou SE, Tsiamadis V, Plevraki E, Petrovic T, et al. Neethling vaccine proved highly effective in controlling lumpy skin disease epidemics in the Balkans. Prev Vet Med 2020; 181:104595.

71. Agianniotaki EI, Babiuk S, Katsoulos PD, Chaintoutis SC, Praxitelous A, Quizon K, et al. Colostrum transfer of neutralizing antibodies against lumpy skin disease virus from vaccinated cows to their calves. Transbound Emerg Dis 2018; 65:2043–8.

72. DAHD. Lumpy Skin Disease, Monthly Report of the Department of Animal Husbandary, Government of Rajasthan, India. 2022.

