## Supplementary Fig S1 for "Evaluation of the safety, immunogenicity and efficacy of a new live-attenuated lumpy skin disease vaccine in India"

### Slide 1
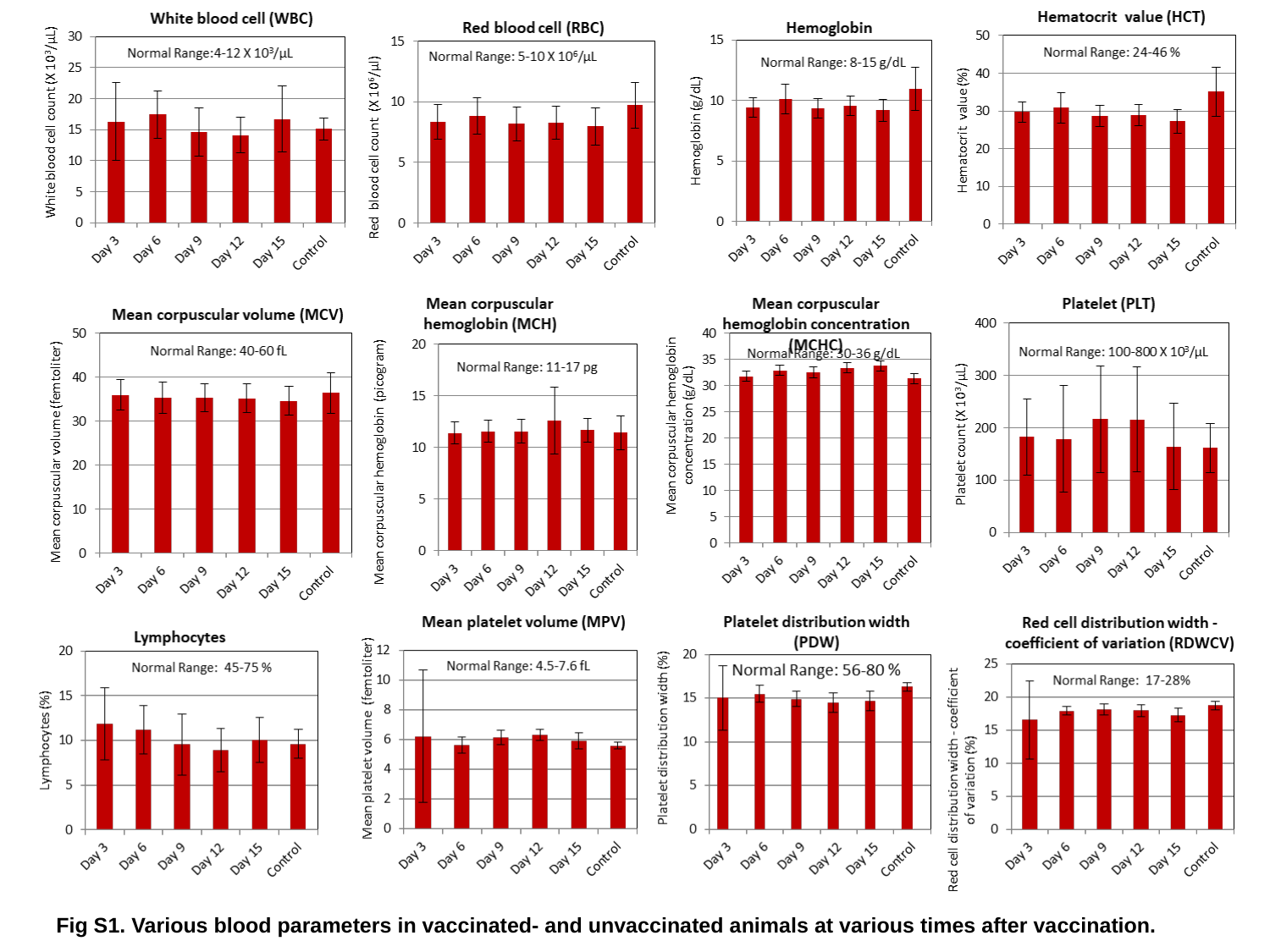

Fig S1. Various blood parameters in vaccinated- and unvaccinated animals at various times after vaccination.
